# Retention dependences support highly confident identification of lipid species in human plasma by reversed-phase UHPLC/MS

**DOI:** 10.1101/2021.05.17.444517

**Authors:** Zuzana Vaňková, Ondřej Peterka, Michaela Chocholoušková, Denise Wolrab, Robert Jirásko, Michal Holčapek

**Affiliations:** University of Pardubice, Faculty of Chemical Technology, Department of Analytical Chemistry, Studentská 573, 53210 Pardubice, Czech Republic

**Keywords:** Lipids, Lipidomics, Ultrahigh-performance liquid chromatography, Reversed-phase, Mass spectrometry, Human plasma, Retention behavior, Fragmentation behavior

## Abstract

Reversed-phase ultrahigh-performance liquid chromatography q mass spectrometry (RP-UHPLC/MS) method was developed with the aim to unambiguously identify a large number of lipid species from multiple lipid classes in human plasma. The optimized RP-UHPLC/MS method employed the C18 column with sub-2 μm particles with the total run time of 25 min. The chromatographic resolution was investigated with 42 standards from 18 lipid classes. The UHPLC system was coupled to high-resolution quadrupole – time-of-flight (QTOF) mass analyzer using electrospray ionization (ESI) measuring full scan and tandem mass spectra (MS/MS) in positive- and negative-ion modes with high mass accuracy. Our identification approach was based on *m/z* values measured with mass accuracy within 5 ppm tolerance in the full scan mode, characteristic fragment ions in MS/MS, and regularity in chromatographic retention dependences for individual lipid species, which provides the highest level of confidence for reported identifications of lipid species including regioisomeric and other isobaric forms. The graphs of dependences of retention times on the carbon number or on the number of double bond(s) in fatty acyl chains were constructed to support the identification of lipid species in homologous lipid series. Our list of identified lipid species is also compared with previous publications investigating human blood samples by various MS based approaches. In total, we have reported more than 500 lipid species representing 26 polar and nonpolar lipid classes detected in NIST Standard reference material 1950 human plasma.

## Introduction

Eukaryotic cells of animal and plant origin contain thousands of lipid species, where energy storage, building blocks, and signaling belong to the main biological functions of these important biologically active substances [1]. Lipidomics is aimed to understand the function of lipids in biological systems. Dysregulated lipids have been reported for various serious diseases, such as diabetes mellitus [2], cancer [3], cardiovascular diseases [4, 5], obesity [6], neurodegenerative disorders [7], *etc.* [8–10]. LIPID MAPS consortium introduced the classification of lipids into eight main categories (fatty acyls, glycerolipids, glycerophospholipids, sphingolipids, sterols, prenols, saccharolipids, and polyketides), where each category contains many classes and subclasses [11].

The lipidomic analysis is mainly performed by mass spectrometry (MS) either with direct infusion (DI) or connected to separation techniques [12], typically with ESI. In DI-MS, the diluted sample extract is directly infused into the mass spectrometer without any chromatographic separation, which provides the advantage for the quantitative analysis due to constant matrix effects, but it lacks the isobaric resolution achievable by chromatography. The identification and quantitation in DI-MS is based on tandem mass spectrometry (MS/MS) using special scanning events, such as precursor ions, neutral loss, multiple reaction monitoring, or data-dependent acquisition of full MS/MS spectra [8, 12, 13]. The methods based on the DI-MS analysis of human plasma typically reported 200 −400 quantified lipid species depending on the particular configuration with low- or high-resolution MS/MS mode [9, 10, 14, 15].

The ultrahigh-performance liquid chromatography (UHPLC) with columns containing sub-2 μm particles enables fast and highly efficient separation. Chromatographic approaches in lipidomics are divided according to the separation mechanisms of lipid species separation represented by reversed-phase (RP) UHPLC and lipid class separation including normal-phase (NP) UHPLC, ultrahigh-performance supercritical fluid chromatography (UHPSFC), and hydrophilic interaction liquid chromatography (HILIC) [12, 16, 17]. The NP-UHPLC/MS can be used for the separation of nonpolar lipid classes [16], while HILIC mode separates individual polar lipid classes according to their polarity and electrostatic interactions [12, 16, 18–20]. UHPSFC is the most recent chromatographic technique used for lipidomic analysis, which separates lipids according to the polarity of the head group of lipids interacting with the polar stationary phase, similarly as in HILIC, but moreover it enables the separation of nonpolar and polar lipids in one analysis during shorter analysis times with higher chromatographic resolution [21, 22]. The quantitative UHPSFC/MS method allows the analysis of over 200 lipid species in less than 6 minutes, which is a suitable technique for high-throughput and comprehensive lipidomic analysis of biological samples [19, 23]. The lipid class separation approaches are preferred for quantitative analysis, because only one internal standard (IS) per lipid class is sufficient for all coeluted lipid species [12, 24]. In RP mode, the retention of lipid species depends on the length of fatty acyl chains and the number and position of the double bonds [19, 23, 25–28]. RP-UHPLC/MS method with two C18 columns in series [24] was applied for the explanation of retention behavior followed by the identification of 346 lipid species in human plasma from 11 classes in 160 min. Long analysis times enable high separation efficiency, but it is not being acceptable for routine measurements. However, newer RP-UHPLC/MS methods were focused on a higher number of identified lipids with the significantly lower run time, which led to greater overlaps of lipid species and complicated identification [4, 25, 26, 29]. Silver-ion high-performance liquid chromatography (HPLC) and chiral HPLC are special chromatographic techniques with different selectivity to a specific part of the molecule, where silver-ion HPLC is based on the separation according to the number and position of double bonds [30, 31] and chiral HPLC is applied for the separation of TG enantiomers [32, 33].

The goal of this work was the development of RP-UHPLC/MS method for the determination of a large number of lipids in human plasma. Full scan and tandem mass spectra were measured using a high-resolution analyzer in both polarity modes of ESI with mass accuracy better than 5 ppm (in MS mode) for confident identification of the individual lipid species. Characteristic fragment ions of individual lipid classes confirmed with representative standards, systematic retention times depending on the polarity, and the verification of *m/z* and in both polarity modes are the tools that lead to the high confidence of lipid species identification. The characteristic ions (adducts, precursor ions, neutral losses, RCOO^-^, *etc.)* have been found for each analyte and the retention behavior of individual lipids is verified by using the second-degree polynomial regression.

## Materials and methods

### Chemicals and standards

Acetonitrile, 2-propanol, methanol, butanol (all HPLC/MS grade), and additives formic acid and ammonium formate were purchased from Thermo Fisher Scientific Inc. (Waltham, MA, USA). Chloroform stabilized by 0.5-1% ethanol (HPLC grade) was purchased from Merck (Darmstadt, Germany). Deionization water was prepared by Milli-Q Reference Water Purification System (Molsheim, France). The lipid standards were purchased from Avanti Polar Lipids (Alabaster, AL, USA), Merck (Darmstadt, Germany) or Nu-Chek Prep (Elysian, MN, USA).

### Sample preparation

The sample of NIST Standard reference material 1950 human plasma (25 μL) was deproteinized with 250 μL of the butanol – methanol mixture (1:1, *v/v)* [4]. The sample was sonicated in an ultrasonic bath at 25 °C for 15 min, centrifuged at 3,000 rpm (886×g) for 10 min at ambient conditions, and purified using 0.25 μm cellulose filter (OlimPeak, Teknokroma) before transferring into the vials with the glass inserts inside for the analysis.

### Lipid standard preparation

A mixture of lipid standards (Table S-1) for the retention behavior monitoring contained triacylglycerols (TG 19:1-19:1-19:1 and TG 15:0-18:1-d7-15:0), diacylglycerols (DG 18:1-18:1, DG 18:1-18:1-d5, DG 15:0-18:1-d7, and DG 12:1-12:1), monoacylglycerols (MG 18:1, MG 18:1-d7, and MG 19:1), phosphatidylcholines (PC 14:0-14:0, PC 15:0-18:1-d7, PC 18:1-18:1, PC 22:1-22:1, and PC 22:0-22:0), lysophosphatidylcholines (LPC 17:0, LPC 18:1, and LPC 18:1-d7), phosphatidylglycerols (PG 14:0-14:0 and PG 18:1-18:1), lysophosphatidylglycerols (LPG 18:1 and LPG 14:0), phosphatidylserines (PS 14:0-14:0 and PS 16:0-18:1), lysophosphatidylserine (LPS 17:1), phosphatidylethanolamines (PE 14:0-14:0, PE 15:0-18:1-d7, and PE 18:1-18:1), lysophosphatidylethanolamines (LPE 18:1 and LPE 14:0), phosphatidylinositol (PI 15:0-18:1-d7), sphingomyelins (SM 18:1/12:0;O2, SM 18:1/18:1;O2, and SM 18:1-d9/18:1;O2), ceramides (Cer 18:1/12:0;O2, Cer 18:1/18:1;O2, Cer 18:1/17:0;O2, and Cer 18:1-d7/18:0;O2), hexosylceramides (GlcCer 18:1/12:0;O2 and GlcCer 18:1/16:0;O2), dihexosylceramide (LacCer 18:1/12:0;O2), cholesteryl ester (CE 16:0 d7), and cholesterol d7 (Chol d7). The mixture was prepared by mixing of aliquots of the stock solutions dissolved in 2-propanol – chloroform mixture (4:1, *v/v).* The standard mix was stored at −80 °C and 100 times diluted just before RP-UHPLC/MS measurements.

### RP-UHPLC/ESI-MS conditions

The lipid separation was performed on a liquid chromatograph Agilent 1290 Infinity series (Agilent Technologies, Waldbronn, Germany). The final RP-UHPLC/MS method used the following conditions: Acquity UPLC BEH C18 VanGuard Pre-column (5 mm × 2.1 mm, 1.7 μm), Acquity UPLC BEH C18 column (150 mm × 2.1 mm, 1.7 μm), flow rate 0.35 mL/min, injection volume 1 μL, column temperature 55 °C, and mobile phase gradient: 0 min – 35 % B, 8 min – 50 % B, 21 – 23 min – 95 % B, and 24 – 25 min – 35 % B. The mobile phase A was a mixture of acetonitrile – water (60/40, *v/v*) and the mobile phase B was the mixture of acetonitrile – 2-propanol (10/90, *v/v*). Both mobile phases contained 0.1 % formic acid and 5 mM ammonium formate (AF).

The analytical experiments were performed on Xevo G2-XS QTOF mass spectrometer (Waters, Milford, MA, USA). The data was acquired in the sensitivity mode using positive-ion and negative-ion modes under the following conditions: the capillary voltage of 3 kV for positive-ion mode and −1.5 kV for negative-ion mode, sampling cone 20 V, source offset 90 V, source temperature 150 °C, desolvation temperature 500 °C, cone gas flow 50 L/h, and the desolvation gas flow 1,000 L/h. The acquisition range was *m/z* 100 – 1500 with a scan time 0.5 s measured in the continuum profile mode for MS, respectively, *m/z* 100 – 1000 with a scan time 0.1 s for MS/MS. Argon as the collision gas at the collision energy of 30 eV was used for MS/MS experiments in both polarity modes. The peptide leucine enkephalin was used as a lock mass for MS experiments.

### Data processing

The data in RP-UHPLC/MS method was acquired using MassLynx software and the raw data file was subjected to noise reduction by Waters Compression Tool. The correction of lock mass was applied for the better mass accuracy and the file was converted from continuum to centroid mode by the Accurate Mass Measure tool in MassLynx. MS/MS spectra were measured without lock mass correction. QuanLynx tool was used for exporting the peak area (tolerance ±15 mDa and retention time ±0.5 min).

## Results and discussion

### RP-UHPLC separation of lipids

The goal of this work was the highly reliable identification of a high number of lipid species in human plasma samples based on the combined information of accurate masses (below 5 ppm), interpreted fragment ions, and homological dependences in the retention behavior of lipid species using RP-UHPLC/MS method. For this purpose, the C18 column with sub-2 μm particles (150 × 2.1 mm, 1.7 μm) was selected based on our previous work [24]. Mobile phase including a mixture of water – acetonitrile – 2-propanol with the addition of additives (ammonium formate and formic acid) for sufficient separation selectivity and better ionization was described by Damen et al. [25]. We modified the gradient elution and investigated the concentration of ammonium formate in a range from 0 to 15 mM. Fig. S-1 shows the peak areas for individual concentrations of AF for lipid standards with short fatty acyl chains and deuterated standards. Other standards have the same trend (not shown). The highest peak areas are obtained without any additives in the mobile phase, but these conditions are not preferable due to altered separation efficiency, worse peak shapes for some polar lipid species, and retention and ionization stability issues leading to irreproducible results. The best compromise in the context of chromatographic separation and ionization efficiency is obtained with 5 mM AF and 0.1 % formic acid (Fig. S-2), which is further used in this work.

A mixture of lipid standards, including deuterated, endogenous, and non-endogenous lipids, was measured in positive-ion and negative-ion ESI modes and used for optimization of the separation process. Fig. S-3 shows the base peak intensity (BPI) chromatograms of 42 lipid species from 18 lipid classes containing LPC, LPE, LPG, LPS, SM, Cer, HexCer, Hex2Cer, PC, PE, PI, PG, PS, MG, DG, TG, Chol, and CE. Lipid species are separated in RP liquid chromatography according to the number of double bonds (DB) and the length of fatty acyl chains (carbon number, CN) [24, 25, 34, 35]. Lipid species within individual lipid classes are retained according to their equivalent carbon number (ECN), which is defined as ECN = CN – 2DB [24, 34]. The higher carbon number in fatty acyl chains corresponds to increased retention times (tR), while an additional DB has the opposite effect and leads to faster elution. Some retention overlap of ECN groups may occur, especially in case of lipid species with high DB number. The nonpolar character of TG and CE leads to the highest retention for these classes, while the more polar lipid classes, such as PC, PE, PI, Cer, glycosylceramides, SM, DG, and Chol, are eluted in a wide retention time range in the middle part of the chromatogram. The most polar lipids (lysoglycerophospholipids, MG, FA, and acylcarnitines (CAR)) are eluted first at the beginning of the chromatogram.

Base peak intensity chromatograms of human plasma sample (Fig. 1) measured by RP-UHPLC/MS in (**a**) positive-ion mode covering 23 lipid classes (LPC, LPC-O, LPC-P, LPE, LPE-P, CAR, MG, PC, PC-P, PC-O, PE, PE-P, monosialodihexosylgangliosides (GM3), SM, Cer, HexCer, dihexosylceramides (Hex2Cer), trihexosylceramides (Hex3Cer), tetrahexosylceramides (Hex4Cer), DG, Chol, CE, and TG) and (**b**) negative-ion modes containing 20 lipid classes (LPC, LPC-O, LPC-P, LPE, LPE-P, lysophosphatidylinositol (LPI), fatty acids (FA), PC, PC-P, PC-O, PE, PE-P, PI, GM3, SM, Cer, HexCer, Hex2Cer, Hex3Cer, and Hex4Cer). MG, DG, TG, CAR, CE, and Chol are detected only in the positive-ion mode, while LPI, PI, and FA are observed only in the negative-ion mode.

**Fig. 1.**
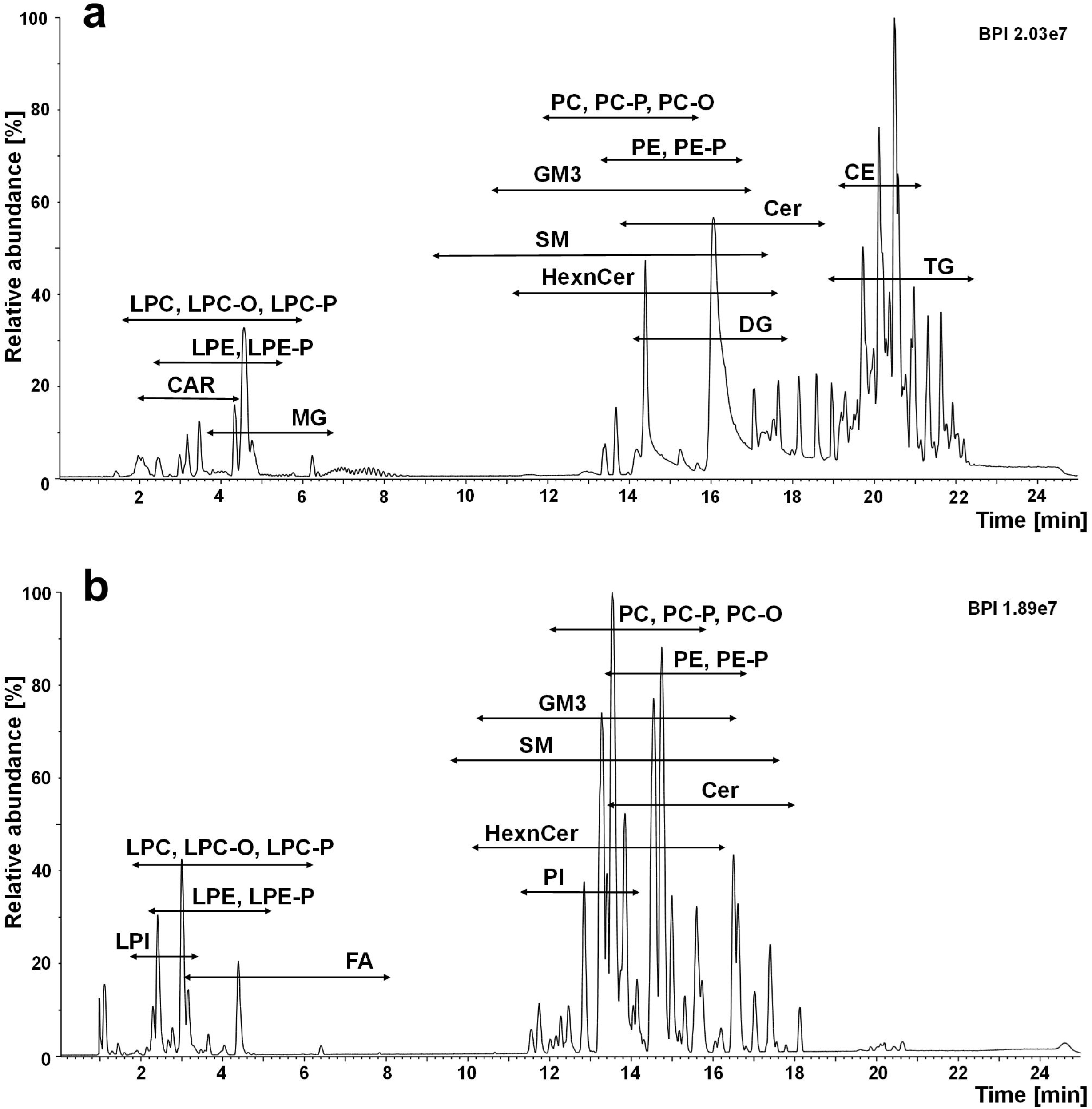
Base peak intensity chromatograms of NIST SRM 1950 human plasma measured by RP-UHPLC/MS in: (**a**) positive-ion and (**b**) negative-ion modes. RP-UHPLC conditions are reported in Materials and methods

### Identification of lipids using RP-UHPLC/MS

The ionization and fragmentation behavior of individual lipid classes is well known from previous works [4, 12, 24, 36–39]. Characteristic precursor and product ions (Table 1) were searched in both positive-ion and negative-ion ESI mass spectra for the identification of lipids in human plasma. The regioisomeric glycerophospholipids can be differentiated based on the different relative intensities of characteristic fragment ions in MS/MS, which allows the identification of the prevailing fatty acyl in *sn-2* position [24]. The *sn-*position of fatty acyls of lysophospholipids is performed based on the retention time, where the lysophospholipids with the fatty acyl in *sn*-2 position elute earlier than with the fatty acyl in *sn*-1 position [24, 37].

In total, 503 lipid species from 26 lipid classes including LPC, LPC-O, LPC-P, LPE, LPE-P, LPI, PC, PC-P, PC-O, PE, PE-P, PI, SM, Cer, HexCer, Hex2Cer, Hex3Cer, Hex4Cer, GM3, CE, Chol, CAR, FA, MG, DG, and TG are identified in the sample of NIST Standard reference material 1950 human plasma (Fig. 2). The list of the identified analytes is summarized in the Electronic Supplementary Materials (ESM) (Table S-2 for positive-ion mode and Table S-3 for the negative-ion mode) with their retention times and characteristic ions in MS and MS/MS modes including mass accuracy. The shorthand notation and nomenclature of lipids follow the updated guidelines by Liebisch *et al.* [40]. Individual lipid species are annotated by their class abbreviation followed by the number of carbon atoms and a sum of DB *(e.g.,* PC 36:2). This species level belongs to the lowest annotation level based on the accurate mass of ions in full-scan mass spectra. Fatty alkyl/acyl level is built on the fragment identification in MS or MS/MS spectra, where the underscore separator is used for unknown *sn-*position (*e.g.*, PC 18:0_18:2). The slash separator means the known preference of *sn-*position (*e.g.*, PC 18:0/18:2). If analytes contain a different type of bond than an ester, the shorthand notation for the ether bond is signed with O and O-alk-1-enyl-bond (in neutral plasmalogen) with P (*e.g.*, PC O-16:0/18:0 and PC P-16:0/18:0). The nomenclature of sphingolipids is based on the presence of sphingosine backbone in the molecule, the number after the shorthand notation reports the CN:DB of the sphingosine base following the number of oxygens and .*N*-linked fatty acyl (*e.g.*, SM 18:1;O2/16:0). The accurate *m/z* value in MS mode (better than 5 ppm), characteristic fragment ions, and neutral losses reflecting the lipid head groups and linked fatty acyls in MS/MS mode, and retention times in RIC chromatograms were considered for the correct identification of lipids. The ions observed in the positive-ion mode include protonated [M+H]^+^, ammoniated [M+NH_4_]^+^, sodiated [M+Na]^+^ molecules, or the loss of water [M+H-H_2_O]^+^. MG, DG, and TG form [M+NH_4_]^+^ and/or [M+Na]^+^ adducts. In case of DG and TG, the [M+H-R_i_COOH]^+^ ions are already found in the full scan spectra, which allows the determination of the fatty acyl composition. CE provide [M+NH_4_]^+^ adducts and the product ion at *m/z* 369 (already in the full scan spectra) belongs to the characteristic head group fragment ion of these compounds containing a cholesterol part. The fragment ion *m/z* 85 belongs among the most abundant fragment ions in MS/MS spectra of CAR. Lipid compounds with the characteristic product ion *m/z* 184 (the cholinephosphate head group) are SM, LPC, LPC-P, LPC-O, PC, PC-O, and PC-P. The loss of ethanolaminephosphate (NL *m/z* 141) is typical for LPE and PE species. MS/MS spectra of PE plasmalogens show two more abundant ions than the neutral loss of 141, *i.e.,* vinyl ether substituent (characteristic fragment ions of *sn-1* fatty acyl position) and acyl substituent (characteristic fragment ions of *sn-2* position fatty acyl). The determination of the composition of sphingolipids (SM, Cer, HexnCer, and GM3) is based on the identification of the type of sphingoid base using the characteristic fragment ions, *i.e.*, *m/z* 236 for 16:1;O2, *m/z* 250 for 17:1;O2, *m/z* 266 for 18:0;O2, *m/z* 264 for 18:1;O2, *m/z* 262 for 18:2;O2, *m/z* 278 for 19:1;O2, and *m/z* 292 for 20:1;O2. The ions observed in the negative-ion MS mode include [M-H]^-^, adducts with formate [M+formate]^-^, or the loss of methyl group [M-CH3]^-^ for choline containing lipid classes. The easiest way to identify fatty acyls is the negative-ion MS mode as illustrated in Fig. 3. In negative-ion MS/MS mode, PC, LPC, and SM form the product ion at *m/z* 168 (choline phosphate head group), LPI and PI provide the fragment ion at *m/z* 241 (inositol phosphate head group) and the neutral loss *Δm/z* 140 (ethanolamine phosphate head group) is typical for LPE, PE, and their plasmalogens or ethers. The identification of the composition of fatty acyls in glycerophospholipid molecules is based on negative-ion MS/MS measurement, where the fragment ions of fatty acyls [R_i_COO]^-^, neutral loss of fatty acyls, and loss of acyl chains in the ketene form can be observed. The fragment ion *m/z* 290 (*N*-acetylneuraminic acid) in the negative-ion MS/MS is typical for GM3 identification.

**Fig. 2.**
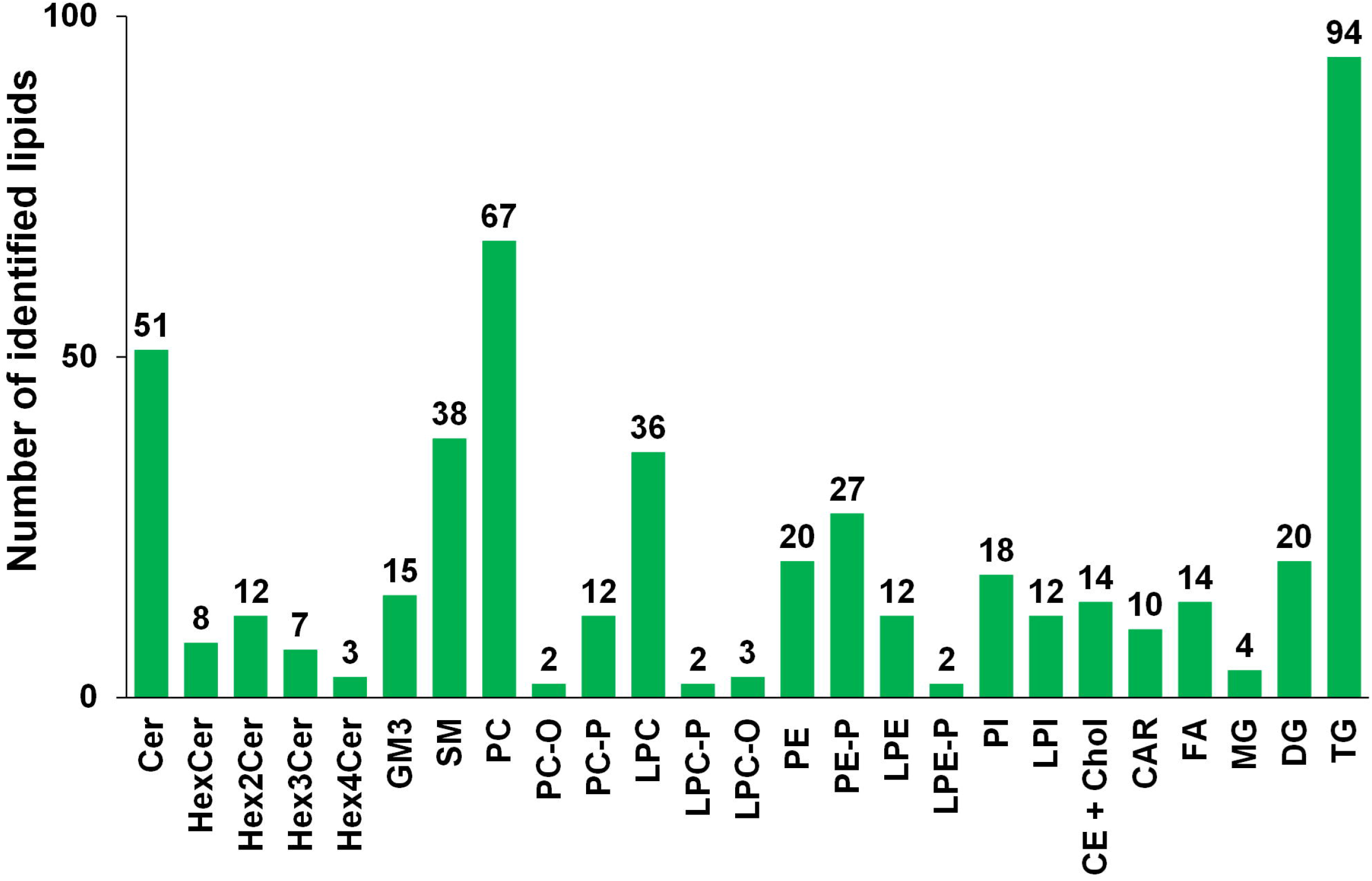
Number of identified lipid species for individual lipid classes in NIST SRM 1950 human plasma

In RP-UHPLC, the retention windows for various lipid classes are overlapped, but the considerable advantage of this chromatographic mode is the separation of isomeric species, for example PC, PC-P, and PC-O (Fig. 3). The identification of these compounds was performed by the retention behavior shown in Table S-4. Furthermore, the chromatographic separation of some isomeric lipid species was achieved, *e.g.*, FA 20:4 (t_R_ 4.22 min and 4.31 min), PC 18:1/20:4 (t_R_ 13.17 min and 13.32 min), PI 18:1/20:4 (t_R_ 11,90 min and 12.12 min), *etc.* There are no detectable differences in their MS/MS spectra. The most likely explanation is the possibility of different DB positions in unsaturated fatty acyls. The verification of this assumption would require authentic lipid standards with defined DB positions.

**Fig. 3.**
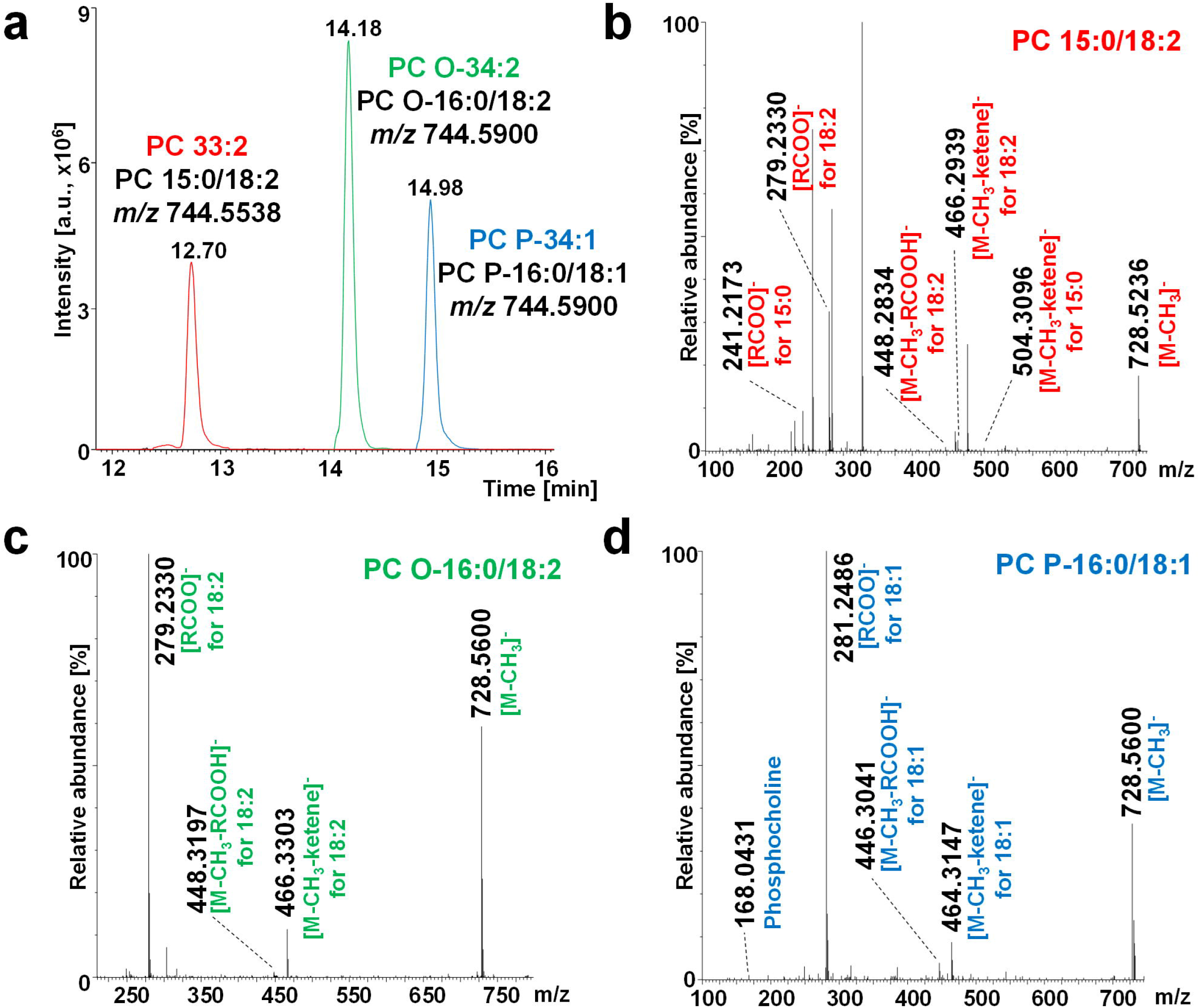
Separation of isobaric phosphatidylcholines and identification of fatty acyl/alkyl position based on characteristic fragment ions annotated in individual MS/MS spectra: (**a**) overlay of reconstructed ion chromatograms, (**b**) MS/MS spectrum of PC 33:2 (PC 15:0/18:2), (**c**) MS/MS spectrum of PC O-34:2 (PC O-16:0/18:2), and (**d**) MS/MS spectrum of PC P-34:1 (PC P-16:0/18:1)

### Retention behavior of various lipid classes

As already mentioned, in RP-UHPLC, the individual lipid species are separated according to the CN and DB. These rules are illustrated in Fig. 4, where part (**a**) depicts the separation according the length of fatty acyl chains in SM species, and part (**b**) the separation of PC species according the number of DB. This retention behavior can be well demonstrated by constructing a dependence of tR on CN or DB by using the second degree of polynomial regression. The typical plots of dependences of tR on the number of the carbon atoms in fatty acyl chains (X) are presented in Fig. 5 for: (**a**) LPC X:0 and LPC X:1, (**b**) CAR X:0, (**c**) SM X:1; O2 and SM X:2;O2, (**d**) Cer X:1; O2 and Cer X:2;O2, (**e**) PC X: 1 and PC X:2, and (**f**) TG X:0 and TG X:2. Fig. 6 summarizes the results of the dependence of tR on the number of DB in fatty acyl chains (Y) for (**a**) LPC 18:Y and LPC 20:Y, (**b**) FA 18:Y, (**c**) DG 36:Y, (**d**) PE P-36:Y and PE P-38:Y, (**e**) PC 34:Y and PC 36:Y, and (**f**) TG 52:Y and 54:Y. Polynomial regression can be also applied for isomers differing only in the regioisomeric positions of fatty acyls. Fig. 7 shows dependences of the t_R_ on the different CN or DB number of lipids for: (**a**) LPC 0:0/18:Y and LPC 18:Y/0:0, (**b**) LPC 0:0/X:0 and LPC X:0/0:0, (**c**) Cer 16:1;O2/X:0 and Cer 18:1;O2/X:0, (**d**) GM3 18:1;O2/X:0 and GM3 18:2;O2/X:0, (**e**) SM 16:1;O2/X:0 and SM 18:1;O2/X:0, and (**f**) PC X:0/18:1 and PC X:0/18:2. Another retention dependences are shown in Fig. S-4 – S-7.

**Fig. 4.**
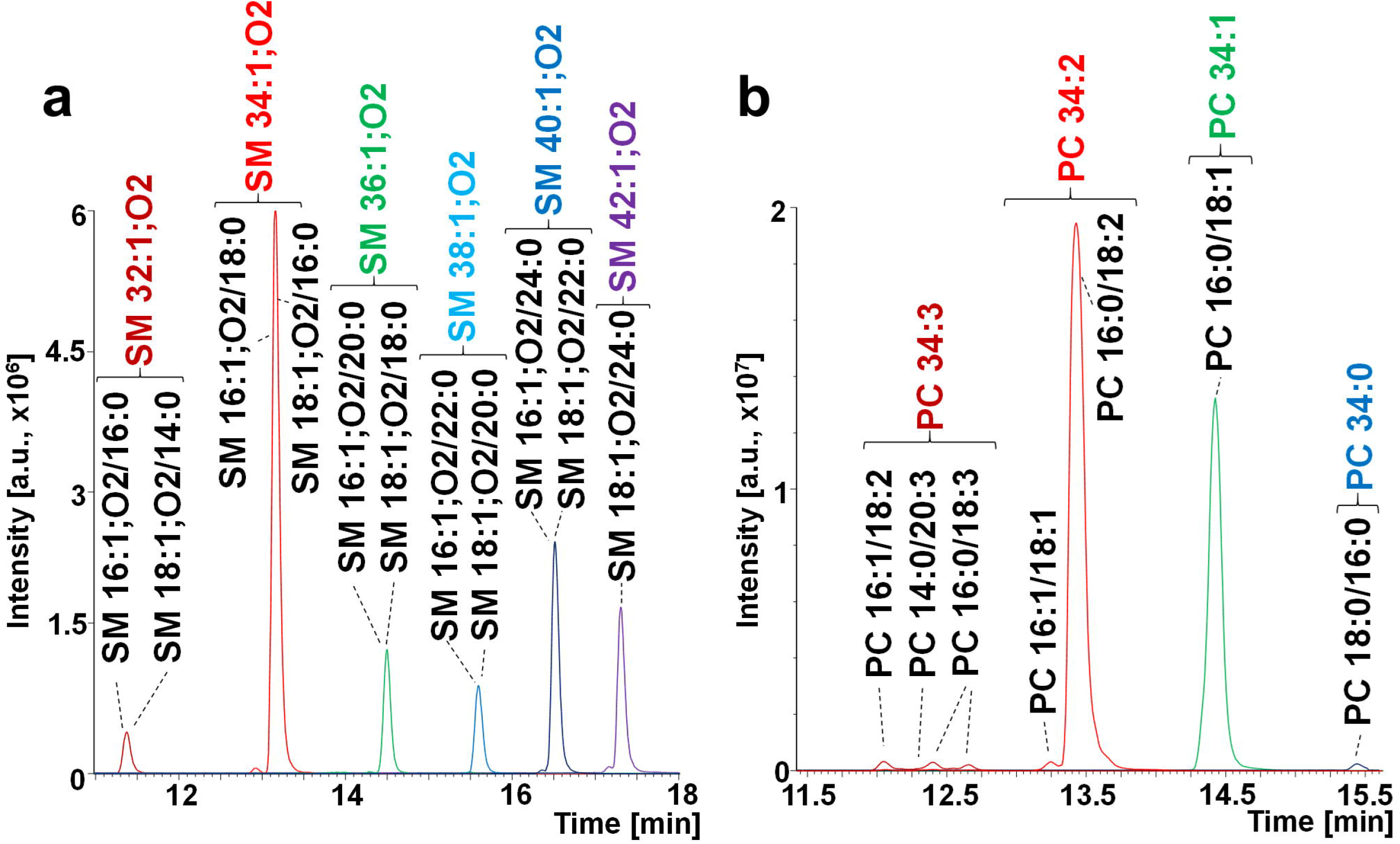
Reconstructed ion chromatograms show the retention behavior of lipid species following a logical series: (**a**) SM differing in the length of fatty acyls and (**b**) PC differing in the number of double bonds

**Fig. 5.**
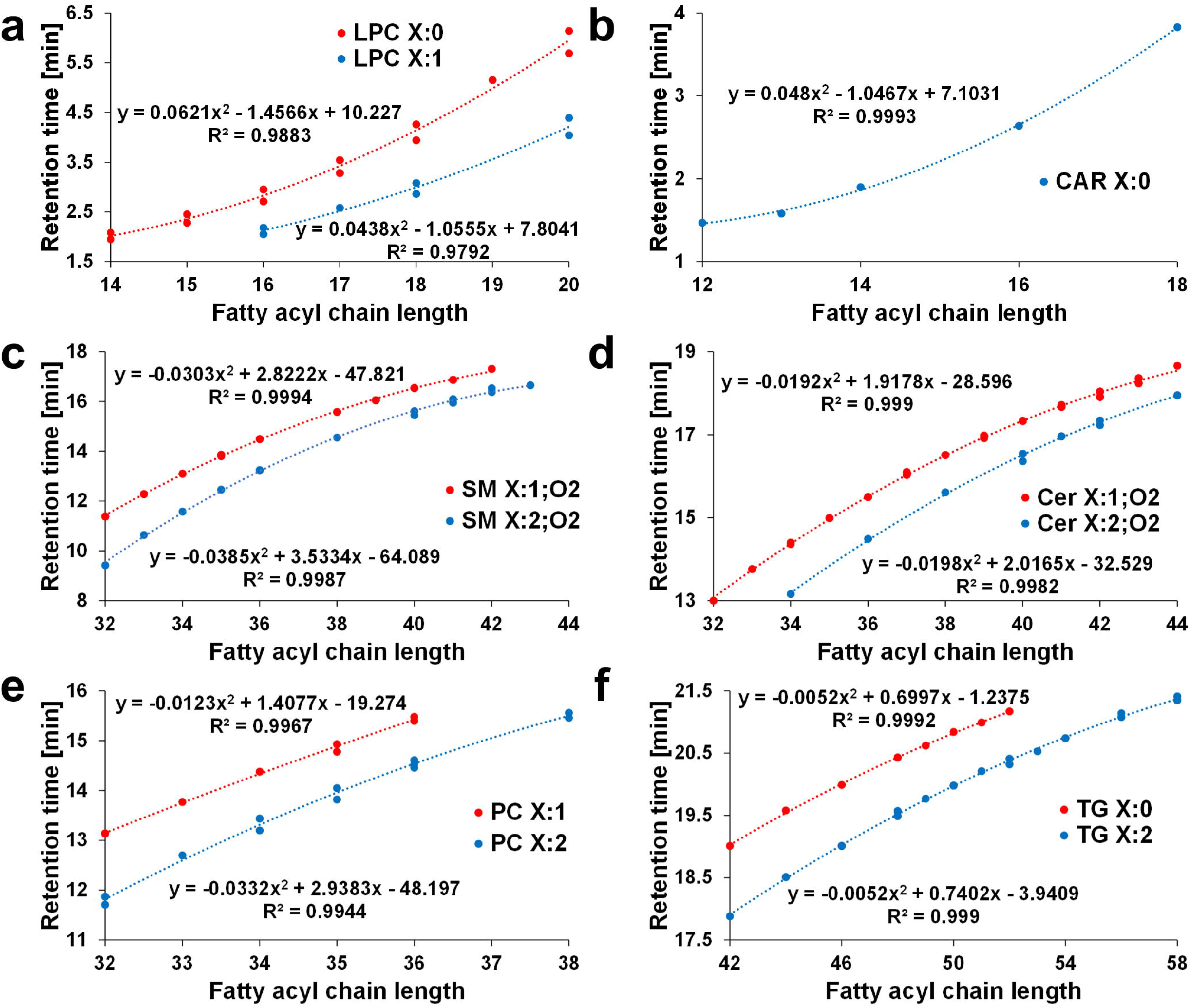
Regular retention behavior of various lipid species illustrated with polynomial dependences of the retention times on the length of fatty acyl chains (X = carbon number): (**a**) LPC X:0 and LPC X:1, (**b**) CAR X:0, (**c**) SM X:1;O2 and SM X:2;O2, (**d**) Cer X:1;O2 and Cer X:2;O2, (**e**) PC X:1 and PC X:2, (**f**) TG X:0 and TG X:2 (X representing number of carbons)

**Fig. 6.**
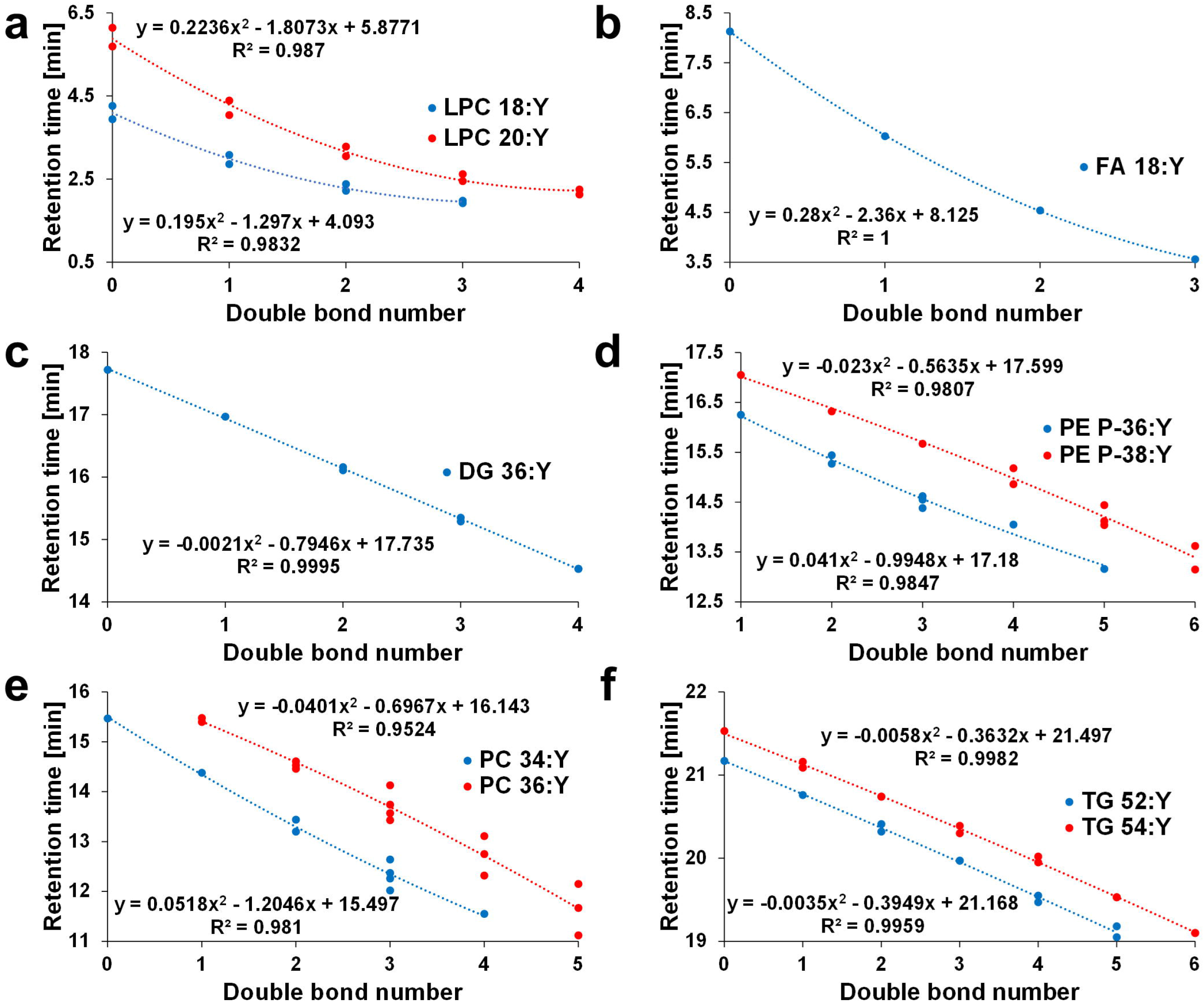
Regular retention behavior of various lipid species illustrated with the polynomial dependences of the retention times on the number of double bond(s) (Y): (**a**) LPC 18:Y and 20:Y, (**b**) FA 18:Y, (**c**) DG 36:Y, (**d**) PE P-36:Y and PE P-38:Y, (**e**) PC 34:Y and PC 36:Y, and (**f**) TG 52:Y and TG 54:Y (Y representing number of double bonds)

**Fig. 7.**
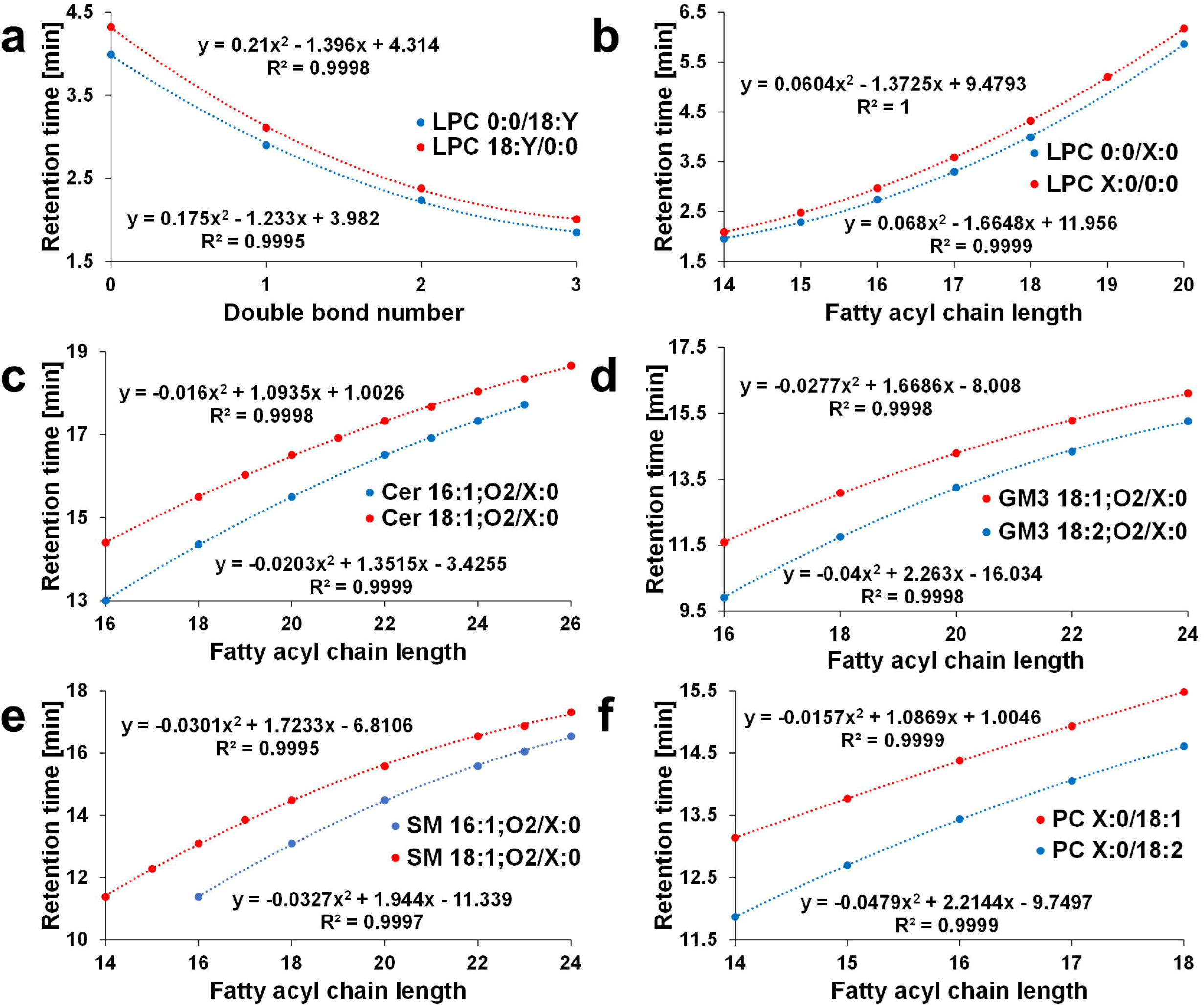
Fatty acyl positions differentiated for various isobaric lipid species. Retention dependences illustrated with the polynomial dependences of retention times on the number of double bonds (X=carbon number in fatty acyls): (**a**) LPC 0:0/18:Y and LPC 18:Y/0:0, and on the length of the fatty acyl chain: (**b**) LPC 0:0/X:0 and PC X:0/0:0, (**c**) Cer 16:1;O2/X:0 and Cer 18:1;O2/X:0, (**d**) GM3 18:1;O2/X:0 and GM3 18:2;O2/X:0, (**e**) SM 16:1;O2/X:0 and SM 18:1;O2/X:0, and (**f**) PC X:0/18:1 and PC X:0/18:2

In many cases, the correlation coefficients R^2^ are better than 0.999, especially for graphs containing differentiated regioisomers, which provides very strong supporting information for the lipid identification in addition to mass spectra. However, R^2^ of plots with unknown composition of fatty acyls may be slightly worse due to the presence of multiple isomers with different t_R_. If some member of the logical CN or DB series is missing, then t_R_ can be predicted from retention dependences with a very high level of accuracy and subsequently the chromatogram may be rechecked for the possible presence of a minor peak at the given t_R_.

### Comparison of new method with the literature

The list of identified lipid species was compared with lipids reported in the literature including lipid species [4, 10, 24, 41, 42] and lipid class [10, 41–43] separation approaches. Comparison of the number of lipid species is shown in ESM Table S-5 and individual lipid species in ESM Table S-6. [4, 10, 24, 41–43]. If compared to our previous RP-UHPLC/MS method [24], we are able to detect more lipid species (503 *vs*. 346 lipid species) and lipid classes (26 *vs.* 16 lipid classes) in much shorter time (25 vs. 160 minutes). In case of TG, dedicated methods based on nonaqueous RP-UHPLC/MS could provide an increased number of identified TG [29], because this generic method is designed for multiple lipid classes favoring the resolution of polar phospholipids and sphingolipids, which results in suboptimal conditions for TG as the least polar lipid class. If we compare the RP-UHPLC/MS method with other methods used in our laboratory for the lipidomic quantitation of biological samples, such as HILIC/MS [43], and UHPSFC/MS [43], the RP-UHPLC provides the highest number of detected lipid species (often annotated at fatty acyl/alkyl level), but this approach requires a larger effort during the identification step and suffers from a lower suitability for quantitative workflows due to the fact that the IS and analytes do not coelute unlike to lipid class separation approaches.

## Conclusions

In this work, we reported 503 lipid species representing 26 lipid classes by RP-UHPLC/MS method using a high confidence level of the identification strategy called 0% false discovery rate in addition to the obvious effort to report the high number of identified lipids. This strategy slightly reduces the number of reported lipids, but we reported only unambiguous identifications with the highest possible confidence supported by multiple criteria to avoid any overreporting in our data set [44]. Some previous attempts to reach the highest possible number resulted in a high level of false identification [45, 46], which is – in our opinion – not the right way for the scientific progress in lipidomics. Therefore, we advocated the strategy based on the highest achievable confidence of identification in parallel with the still relatively high number of reported lipid species and acceptable analysis time for high-throughput analysis. The next step will be the quantitation and comparison of this lipid species separation approach with well-established lipid class separation-based lipidomic quantitation, because the merging of the separation efficiency of RP system together with the quantitation robustness of lipid class separation approaches would be an important step forward in the lipidomic analysis.

## Supporting information

Supplementary information Part 1

Supplementary information Part 2

## Abbreviations

BPI: Base peak intensity
CAR: Acylcarnitine
CE: Cholesteryl ester
Cer: Ceramide
CN: Carbon number
DB: Double bond
DG: Diacylglycerol
DI: Direct infusion
ECN: Equivalent carbon number
ESI: Electrospray ionization
FA: Fatty acid
GlcCer: Glucosylceramide
GM3: Monosialodihexosylganglioside
HexCer: Hexosylceramide
Hex2Cer: Dihexosylceramide
Hex3Cer: Trihexosylceramide
Hex4Cer: Tetrahexosylceramide
HILIC: Hydrophilic interaction liquid chromatography
HPLC: High-performance liquid chromatography
Chol: Cholesterol
IS: Internal standard
LacCer: Lactosylceramide
LPC: Lysophosphatidylcholine
LPE: Lysophosphatidylethanolamine
LPG: Lysophosphatidylglycerol
LPI: Lysophosphatidylinositol
LPS: Lysophosphatidylserine
MG: Monoacylglycerol
MS: Mass spectrometry
MS/MS: Tandem mass spectrometry
NP: Normal-phase
-O: Alkyl bond
-P: Plasmalogen
PC: Phosphatidylcholine
PE: Phosphatidylethanolamine
PG: Phosphatidylglycerol
PI: Phosphatidylinositol
PS: Phosphatidylserine
RIC: Reconstructed ion current
RP: Reversed-phase
SM: Sphingomyelin
TG: Triacylglycerol
tR: Retention time
UHPLC: Ultrahigh-performance liquid chromatography
UHPSFC: Ultrahigh-performance supercritical fluid chromatography
QTOF: Quadrupole – time-of-flight

## Compliance with ethical standards

The ethical approval is not needed for this study, because the biological samples of patients or clinical data are not used in this study.

## References

[1] Wenk MR. The emerging field of lipidomics. Nat Rev Drug Discov. 2005; 4:594–610.

[2] Arneth B, Arneth R, Shams M. Metabolomics of type 1 and type 2 diabetes. Int J Mol Sci. 2019; 20:2467.

[3] Butler LM, Perone Y, Dehairs J, Lupien LE, de Laat V, Talebi A, et al. Lipids and cancer: Emerging roles in pathogenesis, diagnosis and therapeutic intervention. Adv Drug Deliv Rev. 2020; 159:245–93.

[4] Huynh K, Barlow CK, Jayawardana KS, Weir JM, Mellett NA, Cinel M, et al. High-Throughput Plasma Lipidomics: Detailed Mapping of the Associations with Cardiometabolic Risk Factors. Cell Chem Biol. 2019; 26:71–84.

[5] Laaksonen R, Ekroos K, Sysi-Aho M, Hilvo M, Vihervaara T, Kauhanen D, et al. Plasma ceramides predict cardiovascular death in patients with stable coronary artery disease and acute coronary syndromes beyond LDL-cholesterol. Eur. Heart J. 2016; 37:1967–76.

[6] Tonks KT, Coster AC, Christopher MJ, Chaudhuri R, Xu A, Gagnon-Bartsch J, et al. Skeletal muscle and plasma lipidomic signatures of insulin resistance and overweight/obesity in humans. Obesity. 2016; 24:908–16.

[7] van Kruining D, Luo Q, van Echten-Deckert G, Mielke MM, Bowman A, Ellis S, Oliveira TG, Martinez-Martinez P. Sphingolipids as prognostic biomarkers of neurodegeneration, neuroinflammation, and psychiatric diseases and their emerging role in lipidomic investigation methods. Adv Drug Deliv Rev. 2020; 159:232–44.

[8] Wolrab D, Jirásko R, Chocholoušková M, Peterka O, Holcapek M. Oncolipidomics: Mass spectrometric quantitation of lipids in cancer research. TrAC – Trends Anal Chem. 2019; 120:115480.

[9] Wang J, Wang C, Han X. Tutorial on lipidomics. Anal Chim Acta. 2019; 1061:28–41.

[10] Wolrab D, Jirásko R, Cífková E, Höring M, Mei D, Peterka O, et al. Lipidomic profiling of human serum enables detection of pancreatic cancer 1. Nat Com. 2021; revision; preprint available at https://www.medrxiv.org/content/10.1101/2021.01.22.21249767v1

[11] LIPID MAPS. LIPID MAPS Lipidomics Gateway. 2021 (accessed 5.5.2021). https://lipidmaps.org/

[12] Holčapek M, Liebisch G, Ekroos K. Lipidomic Analysis. Anal Chem. 2018; 90:4249–57.

[13] Hsu FF. Mass spectrometry-based shotgun lipidomics – a critical review from the technical point of view. Anal Bioanal Chem. 2018; 410:6387–409.

[14] Sales S, Graessler J, Ciucci S, Al-Atrib R, Vihervaara T, Schuhmann K, et al. Gender, Contraceptives and Individual Metabolic Predisposition Shape a Healthy Plasma Lipidome. Sci Rep. 2016; 6:27710.

[15] Han X, Yang K, Gross RW. Multi-dimensional mass spectrometry-based shotgun lipidomics and novel strategies for lipidomic analyses. Mass Spectrom Rev. 2012; 31:134–78.

[16] Čajka T, Fiehn O. Comprehensive analysis of lipids in biological systems by liquid chromatography–mass spectrometry. TrAC – Trends Anal Chem. 2014; 61:192–206.

[17] Triebl A, Hartler J, Trötzmüller M, C. Köfeler H. Lipidomics: Prospects from a technological perspective. Biochim Biophys Acta – Mol Cell Biol Lipids. 2017; 1862:740–6.

[18] Cífková E, Holcapek M, Lísa M, Vrána D, Melichar B, Študent V. Lipidomic differentiation between human kidney tumors and surrounding normal tissues using HILIC-HPLC/ESI-MS and multivariate data analysis. J Chromatogr B. 2015; 1000:14–21.

[19] Wolrab D, Chocholoušková M, Jirásko R, Peterka O, Holčapek M. Validation of lipidomic analysis of human plasma and serum by supercritical fluid chromatography–mass spectrometry and hydrophilic interaction liquid chromatography–mass spectrometry. Anal Bioanall Chem. 2020; 412:75–2388.

[20] Rampler E, Schoeny H, Mitic BM, el Abiead Y, Schwaiger M, Koellensperger G. Simultaneous non-polar and polar lipid analysis by on-line combination of HILIC, RP and high resolution MS. Analyst. 2018; 143:1250–8.

[21] Dispas A, Jambo H, André S, Tyteca E, Hubert P. Supercritical fluid chromatography: A promising alternative to current bioanalytical techniques. Bioanalysis. 2018; 10:107–24.

[22] Lísa M, Holcapek M. High-Throughput and Comprehensive Lipidomic Analysis Using Ultrahigh-Performance Supercritical Fluid Chromatography-Mass Spectrometry. Anal Chem. 2015; 87:7187–95.

[23] Chocholoušková M, Jirásko R, Vrána D, Gatěk J, Melichar B, Holcapek M. Reversed phase UHPLC/ESI-MS determination of oxylipins in human plasma: a case study of female breast cancer. Anal Bioanal Chem. 2019; 411:1239–51.

[24] Ovčačíková M, Lísa M, Cífková E, Holcapek M. Retention behavior of lipids in reversed-phase ultrahigh-performance liquid chromatography-electrospray ionization mass spectrometry. J Chromatogr A. 2016; 1450:76–85.

[25] Damen CWN, Isaac G, Langridge J, Hankemeier T, Vreeken RJ. Enhanced lipid isomer separation in human plasma using reversed-phase UPLC with ion-mobility/high-resolution MS detection. J Lipid Res. 2014; 55:1772–83.

[26] Yamada T, Uchikata T, Sakamoto S, Yokoi Y, Fukusaki E, Bamba T. Development of a lipid profiling system using reverse-phase liquid chromatography coupled to high-resolution mass spectrometry with rapid polarity switching and an automated lipid identification software. J Chromatogr A. 2013; 1292:211–8.

[27] Fauland A, Köfeler H, Trötzmüller M, Knopf A, Hartler J, Eberl A, Chitraju C, Lankmayr E, Spener F. A comprehensive method for lipid profiling by liquid chromatography-ion cyclotron resonance mass spectrometry. J Lipid Res. 2011; 52:2314–22.

[28] Sandra K, Pereira AS, Vanhoenacker G, David F, Sandra P. Comprehensive blood plasma lipidomics by liquid chromatography/quadrupole time-of-flight mass spectrometry. J Chromatogr A. 1217:4087–99.

[29] Lísa M, Holcapek M. Triacylglycerols profiling in plant oils important in food industry, dietetics and cosmetics using high-performance liquid chromatography-atmospheric pressure chemical ionization mass spectrometry. J Chromatogr A. 2008; 1198-1199:115–30.

[30] Nikolova-Damyanova B. Retention of lipids in silver ion high-performance liquid chromatography: Facts and assumptions. J Chromatogr A. 2009; 1216:1815–24.

[31] Lísa M, Denev R, Holcapek M (2013) Retention behavior of isomeric triacylglycerols in silver-ion HPLC: Effects of mobile phase composition and temperature. J Sep Sci. 2013; 36:2888–900.

[32] Seon HL, Williams M v., Blair IA. Targeted chiral lipidomics analysis. Prostaglandins Other Lipid Mediat. 2005; 77:141–57.

[33] Lísa M, Holcapek M. Characterization of triacylglycerol enantiomers using chiral HPLC/APCI-MS and synthesis of enantiomeric triacylglycerols. Anal Chem. 2013; 85:1852–9.

[34] Holcapek M, Jandera P, Fischer J, Prokeš B. Analytical monitoring of the production of biodiesel by high-performance liquid chromatography with various detection methods. J Chromatogr A. 1999; 858:13–31.

[35] Holcapek M, Ovčačíková M, Lísa M, Cífková E, Hájek T. Continuous comprehensive two-dimensional liquid chromatography–electrospray ionization mass spectrometry of complex lipidomic samples. Anal Bioanal Chem. 2015; 407:5033–43.

[36] Murphy RC Tandem Mass Spectrometry of Lipids: Molecular Analysis of Complex Lipids. 1st ed. Cambridge; Royal Society of Chemistry; 2014.

[37] Lísa M, Cífková E, Holcapek M. Lipidomic profiling of biological tissues using off-line two-dimensional high-performance liquid chromatography-mass spectrometry. J Chromatogr A. 2011; 1218:5146–56.

[38] Hořejší K, Jirásko R, Chocholoušková M, Wolrab D, Kahoun D, Holcapek M. Comprehensive Identification of Glycosphingolipids in Human Plasma Using Hydrophilic Interaction Liquid Chromatography-Electrospray Ionization Mass Spectrometry. Metabolites. 2021; 11:140–63.

[39] Merrill AH, Sullards MC, Allegood JC, Kelly S, Wang E. Sphingolipidomics: High-throughput, structure-specific, and quantitative analysis of sphingolipids by liquid chromatography tandem mass spectrometry. Methods. 2005; 36:207–24.

[40] Liebisch G, Fahy E, Aoki J, Dennis EA, Durand T, Ejsing CS, et al. Update on LIPID MAPS classification, nomenclature, and shorthand notation for MS-derived lipid structures. J Lipid Res. 2020; 61:1539–55.

[41] Quehenberger O, Armando AM, Brown AH, Milne SB, Myers DS, Merrill AH, et al. Lipidomics reveals a remarkable diversity of lipids in human plasma1. J Lipid Res. 2010; 51:3299–305.

[42] Bowden JA, Heckert A, Ulmer CZ, Jones CM, Koelmel JP, Abdullah L, et al. Harmonizing lipidomics: NIST interlaboratory comparison exercise for lipidomics using SRM 1950-Metabolites in frozen human plasma. J Lipid Res. 2017; 58:2275–88.

[43] Wolrab D, Chocholoušková M, Jirásko R, Peterka O, Mužáková V, Študentová H, Melichar B, Holcapek M. Determination of one year stability of lipid plasma profile and comparison of blood collection tubes using UHPSFC/MS and HILIC-UHPLC/MS. Anal Chim Acta. 2020; 1137:74–84.

[44] Liebisch G, Ahrends R, Arita M, Arita M, Bowden JA, Ejsing CS, et al. Lipidomics needs more standardization. Nat Metab. 2019; 1:745–7.

[45] Liebisch G, Ejsing CS, Ekroos K. Identification and annotation of lipid species in metabolomics studies need improvement. Clin Chem. 2015; 61:1542–4.

[46] Kofeler HC, Eichmann TO, Ahrends R, Bowden JA, Danne-Rasche N, Fedorova M, et al. Nat Com. 2021; revision; preprint available at https://zenodo.org/record/4672232#.YJ5c0odxeUl

