## Supplementary information Part 1 for "Retention dependences support highly confident identification of lipid species in human plasma by reversed-phase UHPLC/MS"

**Figures: 7**

**Fig. S-1** Comparison of peak areas *vs.* molar concentrations of ammonium formate (AF) for some selected deuterated and short fatty acyl chain lipid standards, whereas other standards follow the same trends. PI 33:1 d7 and PS 28:0 are not ionized in the mobile phase without additives. Data are presented as the mean value ± standard deviation (SD) from three measurements.

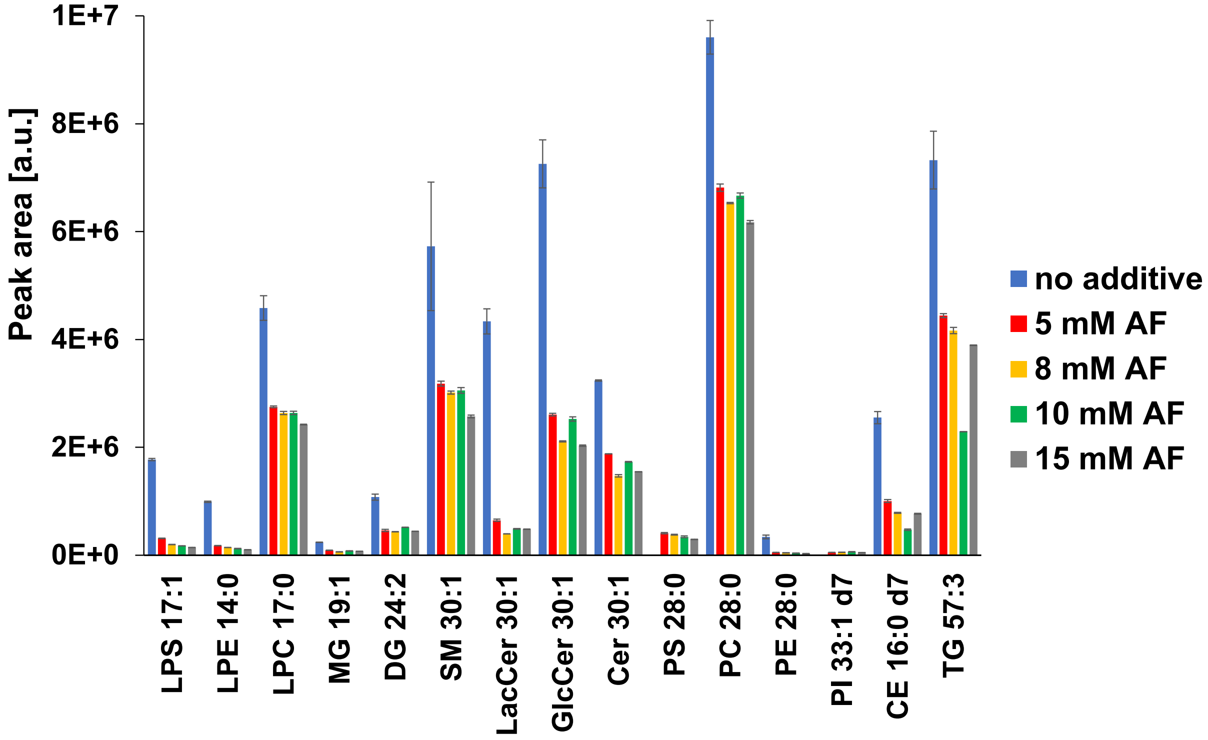

**Fig. S-2** Influence of molar concentration of ammonium formate (AF) on RP-UHPLC separation of lipid standards: (**a**) 0 mM of AF, (**b**) 5 mM AF, (**c**) 8 mM AF, (**d**) 10 mM AF, and (**e**) 15 mM AF in mobile phase. The conditions of analysis are reported in Materials and methods.

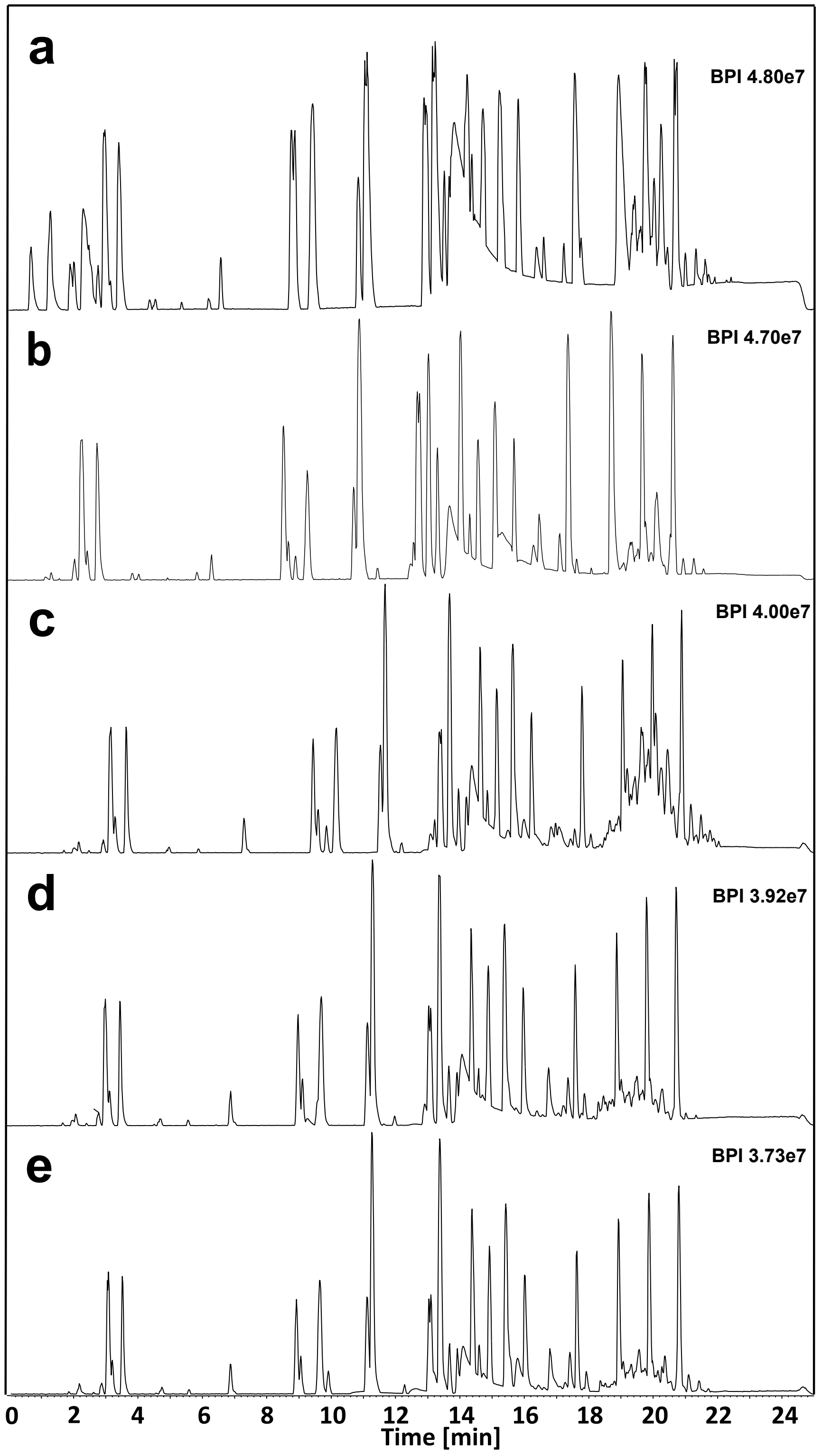

**Fig. S-3** Base peak chromatograms of the lipid standard mixture measured by RP-UHPLC/MS in: (**a**) positive-ion mode and (**b**) negative-ion mode. The conditions of analysis are reported in Materials and methods.

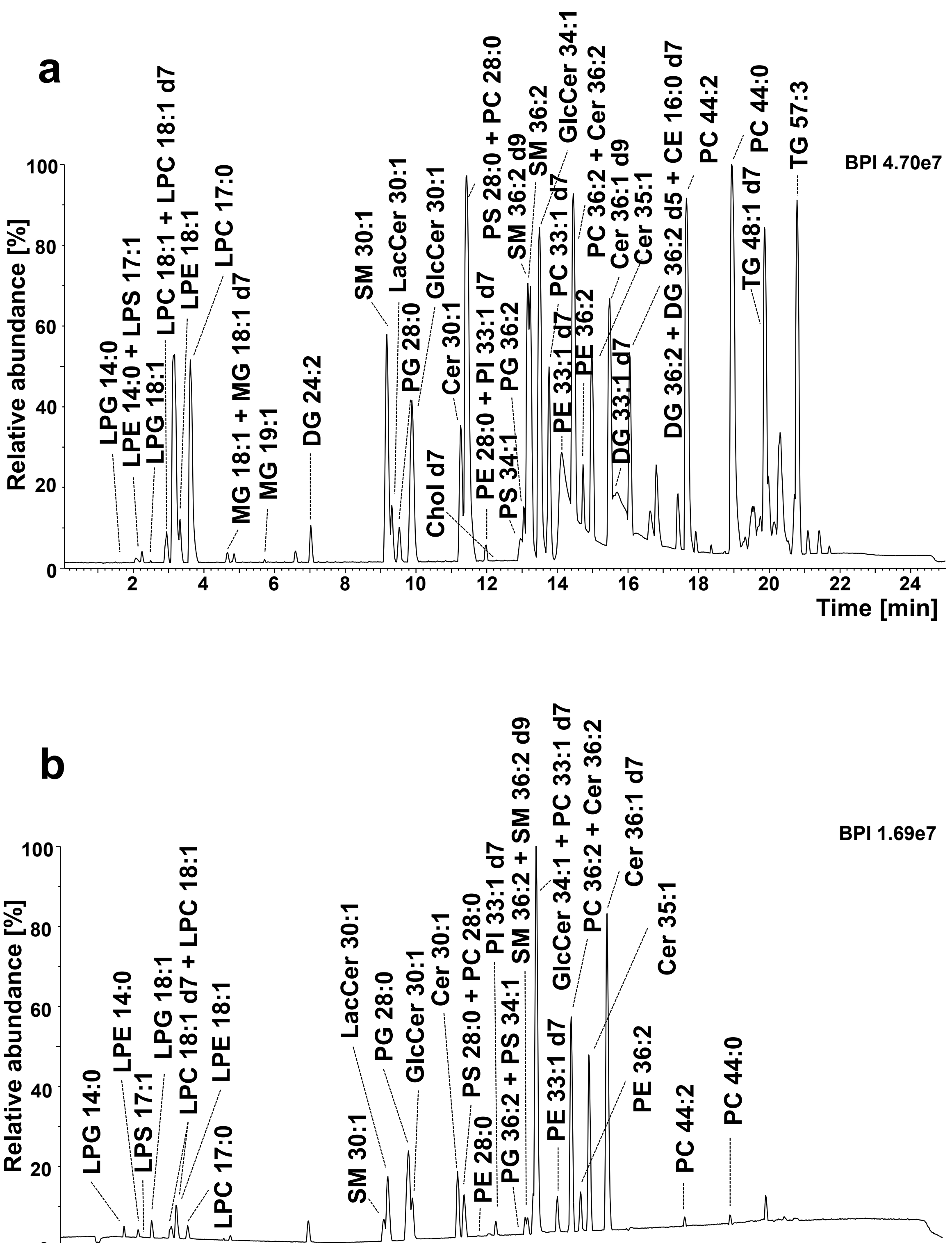

**Fig. S-4** Retention behavior of various lipid species within lipid class shows the polynomial dependences of retention times on the length of fatty acyl chains: (**a**) GM3 X:1;O2 and GM3 X:2;O2, (**b**) PC X:3 and PC X:4, (**c**) TG X:1 and TG X:3, where X represents carbon number and number of present double bonds is written behind colon.

**a**

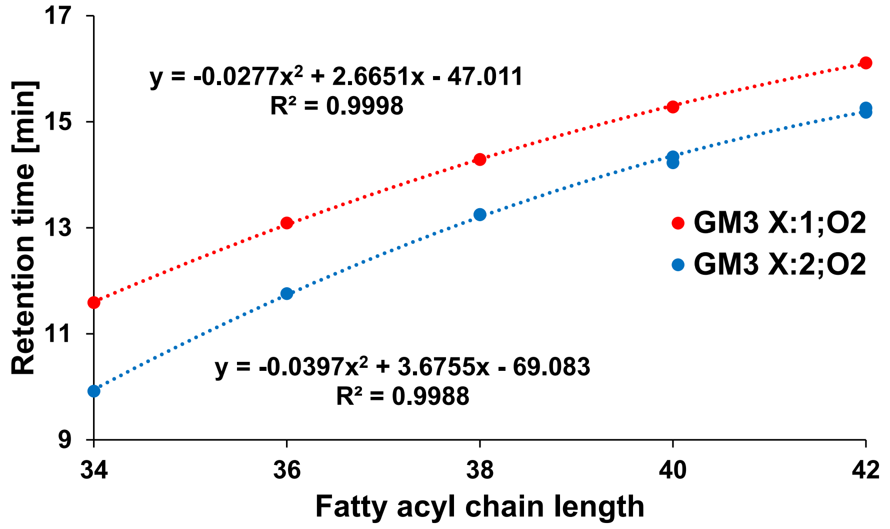


**b**

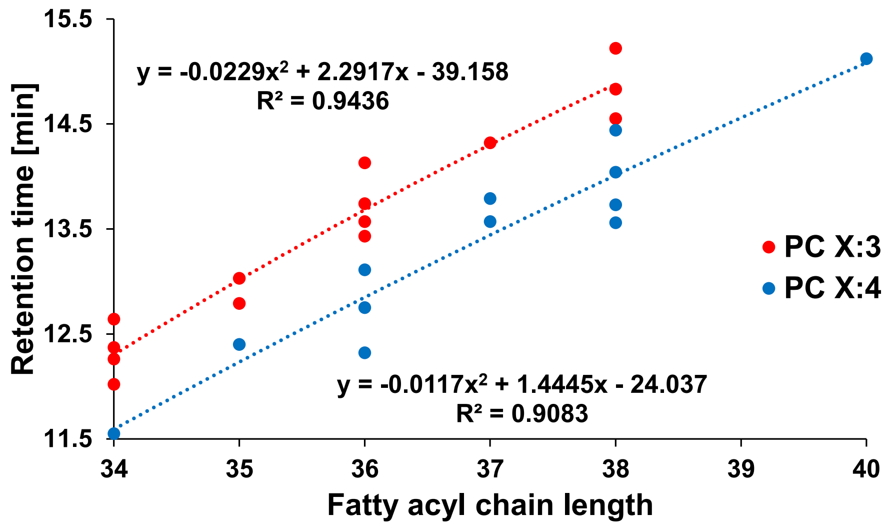

**Fig. S-4** (Continuation)

**c**

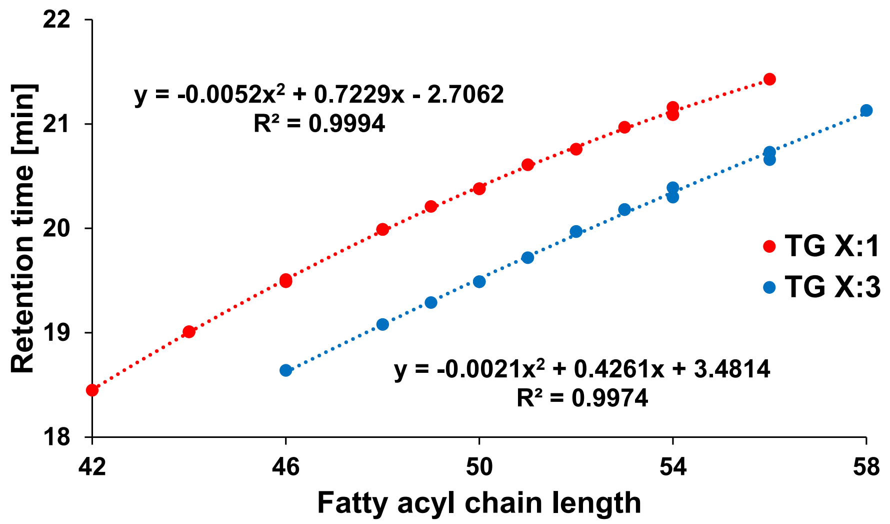

**Fig. S-5** Retention behavior of various lipid species within lipid class shows the polynomial dependences of retention times on the number of double bonds: (**a**) PC 38:Y and PC 40:Y, (**b**) PE 36:Y, PE 38:Y, and PE 40:Y, (**c**) PI 36:Y and PI 38:Y, (**d**) CE 18:Y and CE 20:Y, (**e**) TG 46:X, TG 48:X, and TG 49:X, (**f**) TG 50:Y and TG 51:Y, (**g**) TG 56:Y, where Y represents number of double bond(s) and carbon number is written before colon.

**a**

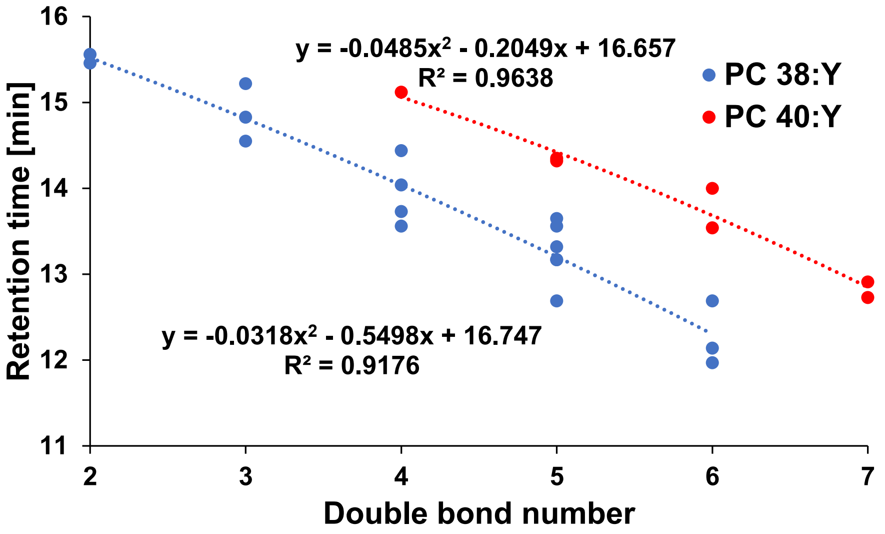


**b**

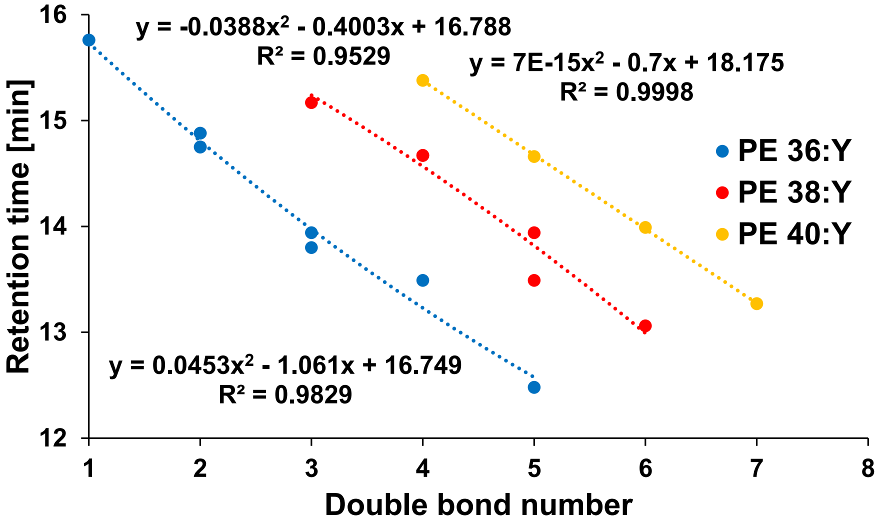

**Fig. S-5** (Continuation)

**c**

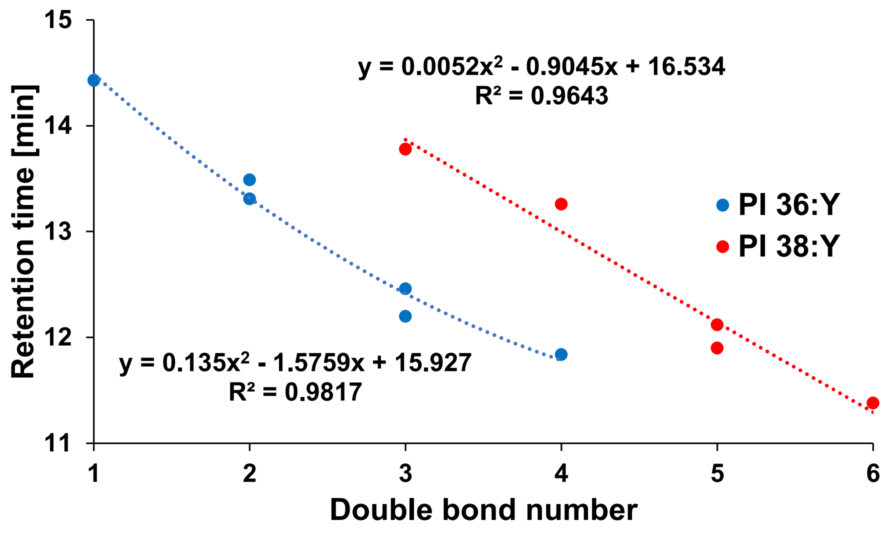


**d**

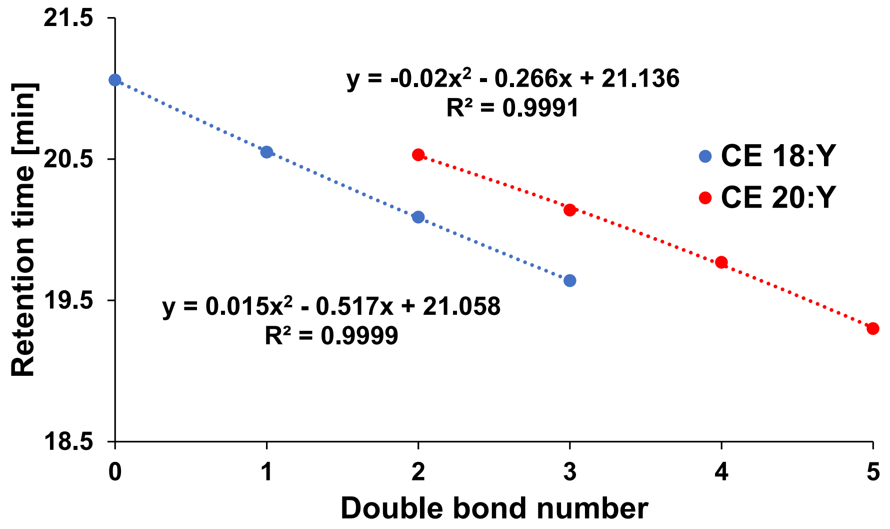

**Fig. S-5** (Continuation)

e

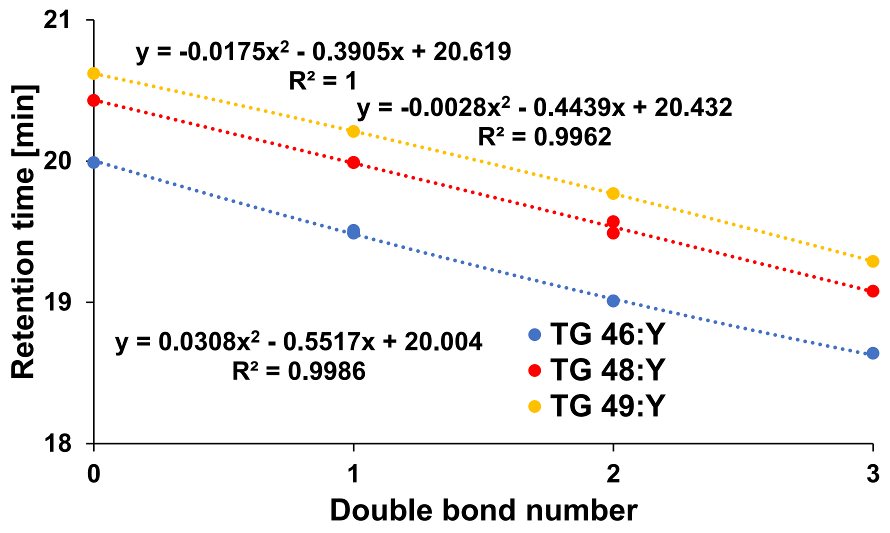


f

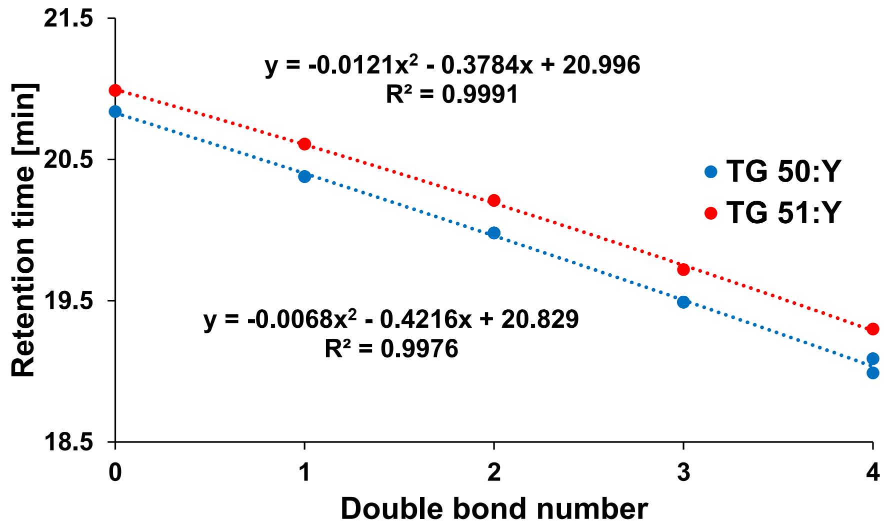

g

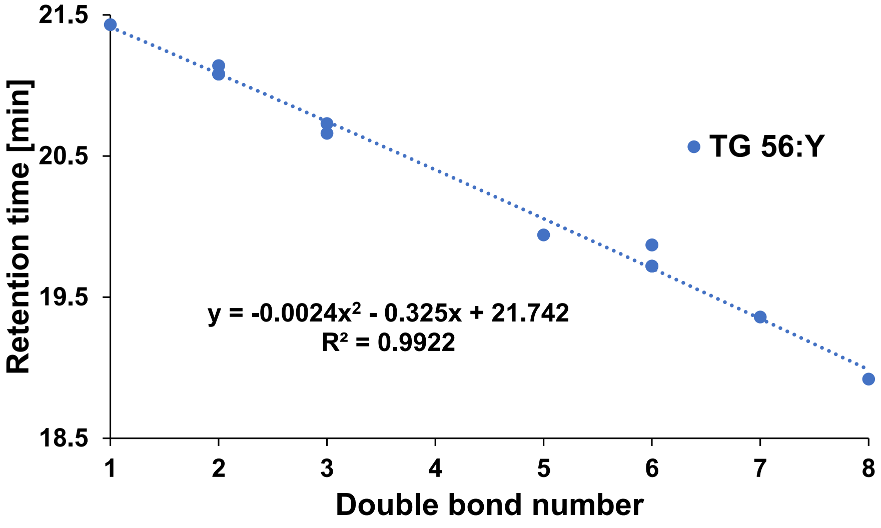

21.5
20.5
19.5
18.5
Retention time [min]
TG 56:Y
y = -0.0024x² - 0.325x + 21.742
R² = 0.9922
1
2
3
4
5
6
7
8
Double bond number

**Fig. S-6** Retention behavior of various lipid species bonding the fatty acyl on the defined *sn*-position or containing the characteristic sphingoid backbone shows the polynomial dependences of retention times on the length of other fatty acyl chains: (**a**) LPC 0:0/X:1, (**b**) PC X:0/20:3 and PC X:0/20:4, (**c**) Cer 18:1;O2/X:0 and Cer 18:2;O2/X:0, (**d**) Cer 17:1;O2/X:0 and Cer 19:1;O2/X:0, (**e**) HexCer 18:1;O2/X:0, Hex2Cer 18:1;O2/X:0, and Hex3Cer 18:1;O2/X:0, (**f**) SM 18:1;O2/X:0 and SM 18:2;O2/X:0, where X represents carbon number and number of present double bonds is written behind colon.

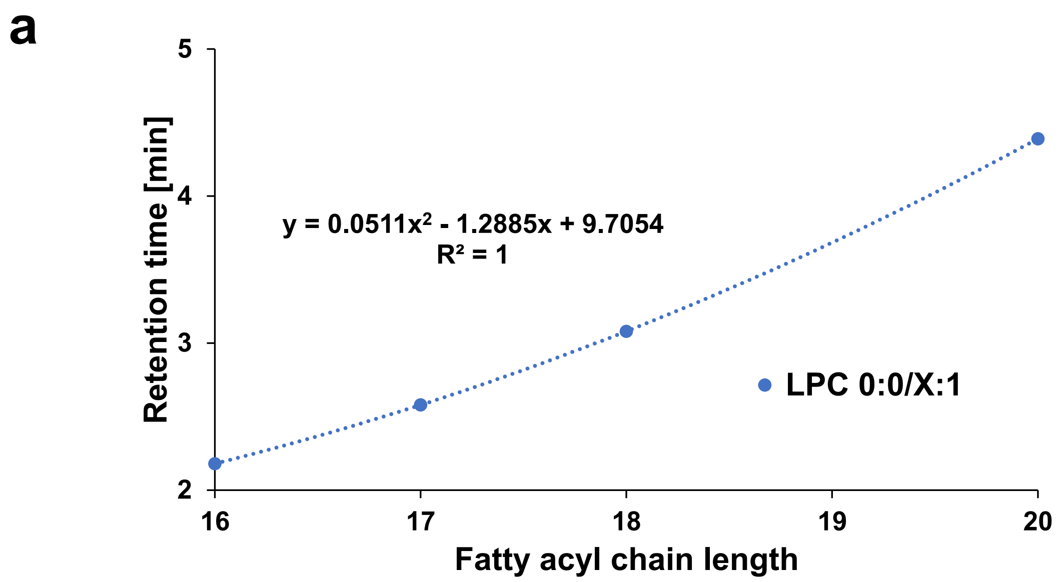


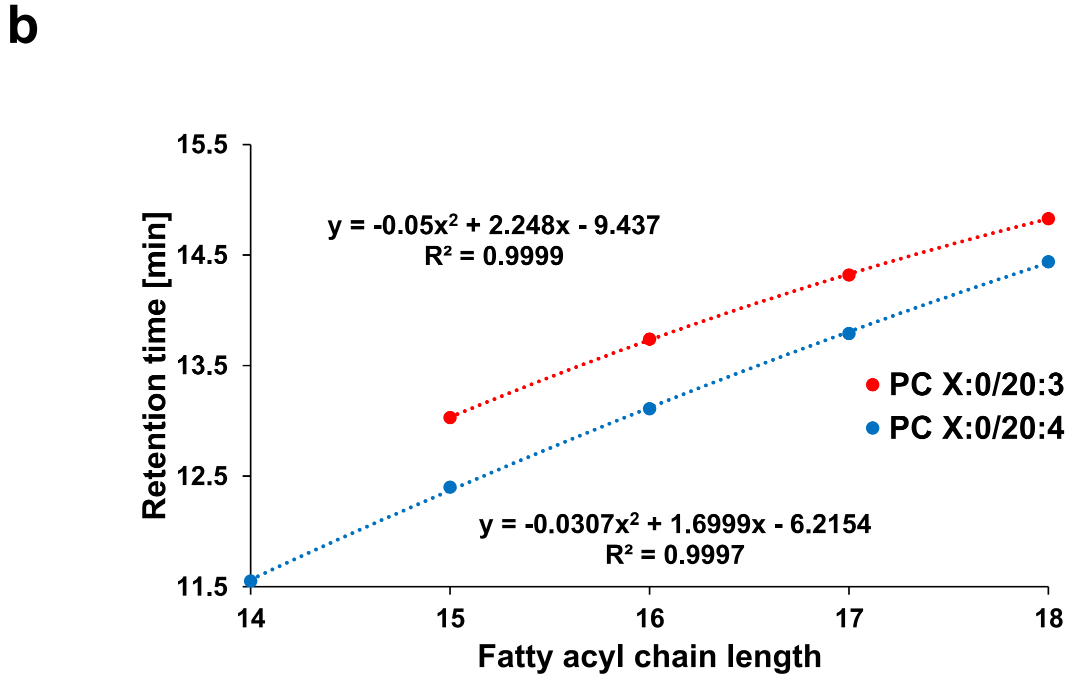

**Fig. S-6** (Continuation)

**c**

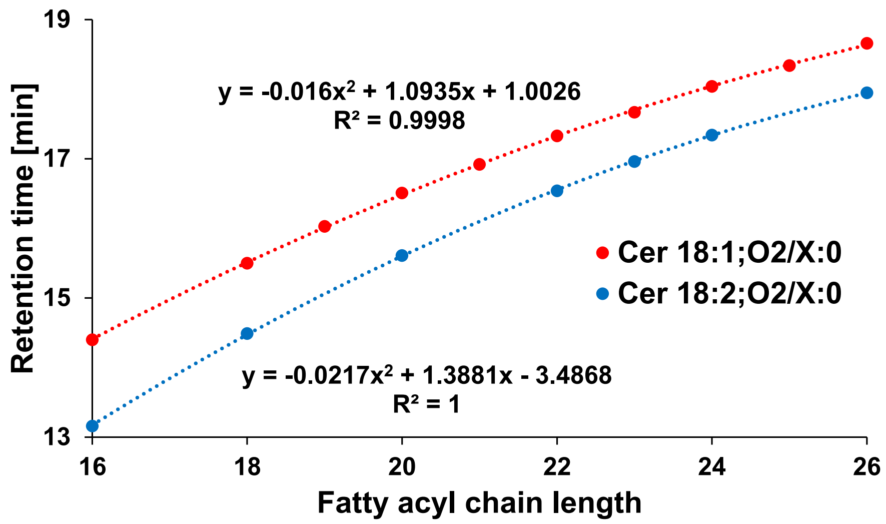


**d**

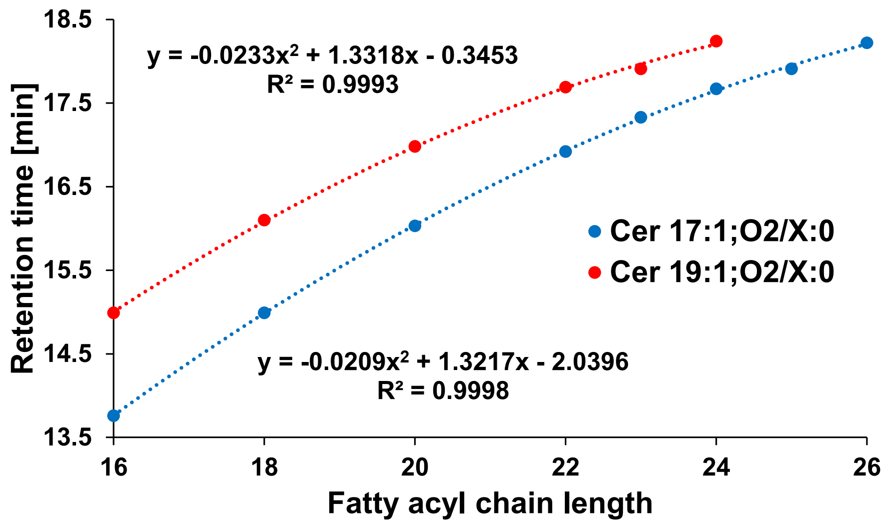

**Fig. S-6** (Continuation)

e

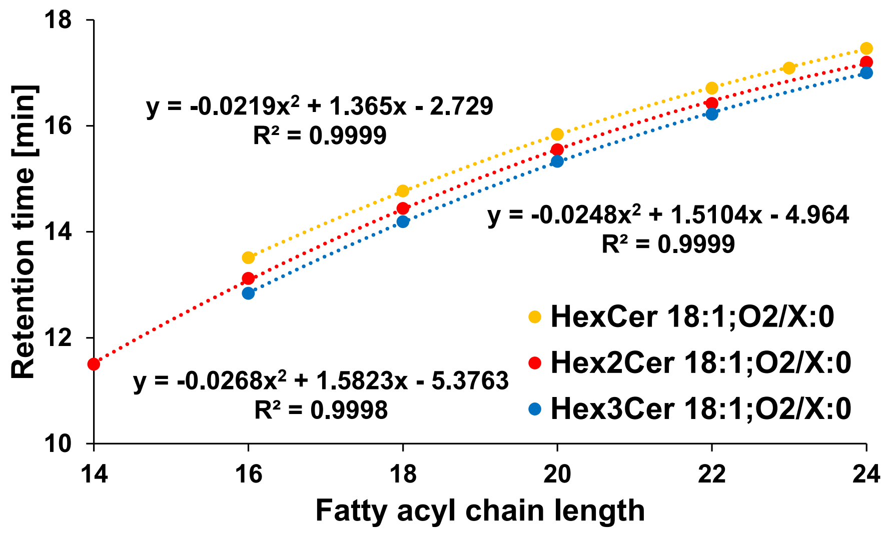


f

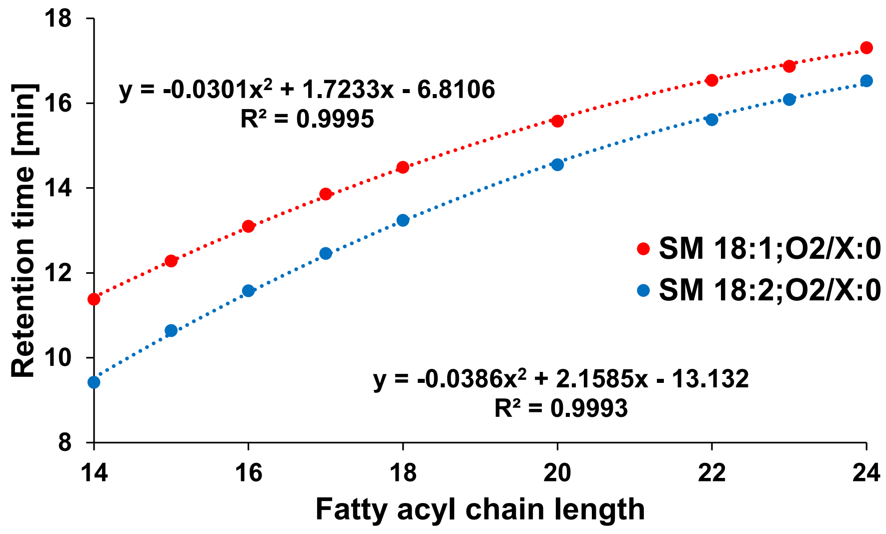

**Fig. S-7** Retention behavior of various lipid species bonding the fatty acyl on the defined *sn*-position shows the polynomial dependences of retention times on the number of double bonds: (**a**) LPC 0:0/20:Y and LPC 20:Y/0:0, (**b**) PC 16:0/20:Y and PC 18:0/20:Y, where Y represents number of double bond(s) and carbon number is written before colon.

a

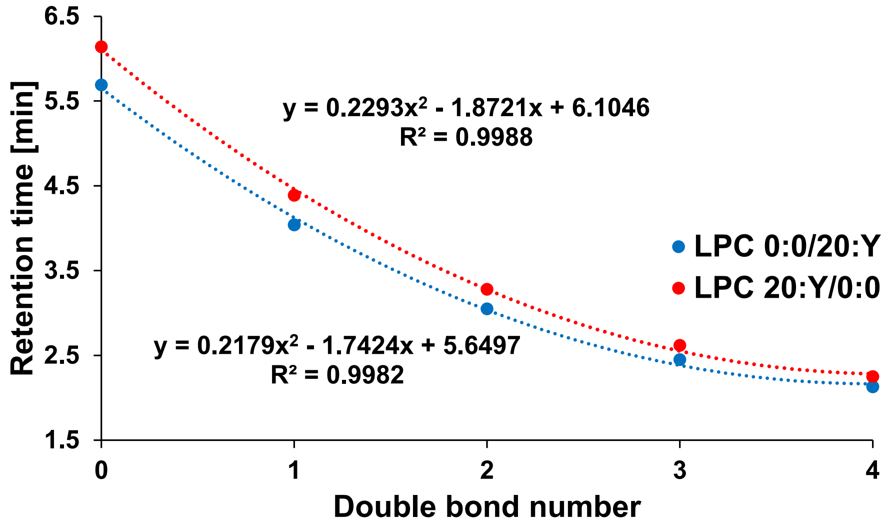


b

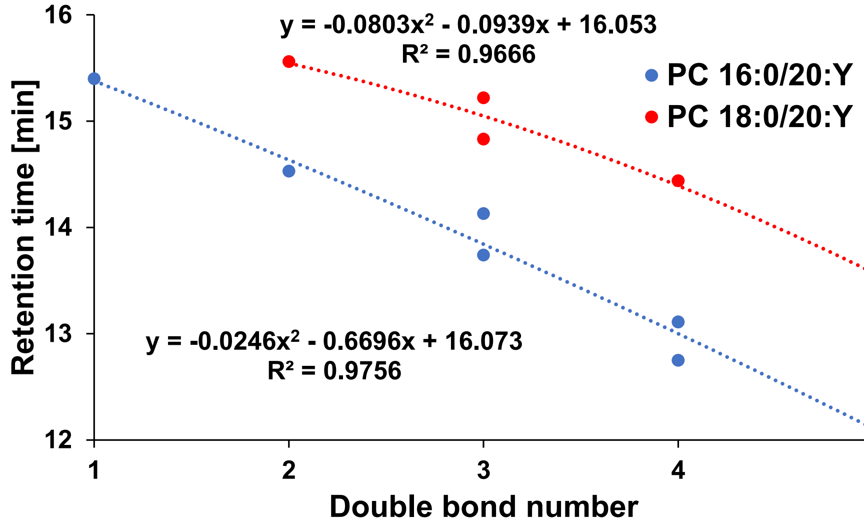
