## Supplementary material for "Retention dependences support highly confident identification of lipid species in human plasma by reversed-phase UHPLC/MS": Table 1

**Table 1** Characteristic ions of individual lipid classes used for the identification of lipid species observed for NIST SRM 1950 human plasma in positive-ion and negative-ion modes by RP-UHPLC/ESI-MS

| **Lipid class** | **Positive-ion mode** | **Negative-ion mode** |
| --- | --- | --- |
| **Cer** | [M+H-H_2_O]^+^, N^II*^ | [M+formate]^-^, P-ion^*^, U-ion^*^, R-ion^*^, T-ion^*^,  S-ion^*^ |
| **HexCer** | [M+H]^+^, N^II*^, Z_0_^*^ | [M+formate]^-^, Z_0_^*^ |
| **Hex2Cer** | [M+H]^+^, N^II*^, Z_0_^*^, Z_1_^*^ | [M+formate]^-^, Z_0_^*^, Z_1_^*^ |
| **Hex3Cer** | [M+H]^+^, N^II*^, Z_0_^*^, Z_1_^*^, Z_2_^*^ | [M+formate]^-^, Z_0_^*^, Z_1_^*^, Z_2_^*^ |
| **Hex4Cer** | [M+H]^+^, N^II*^, Z_0_^*^, Z_1_^*^, Z_2_^*^, Z_3_^*^ | n.d. |
| **GM3** | [M+H]^+^, N^II*^, Z_0_^*^, NeuAc | [M-H]^-^, NeuAc |
| **SM** | [M+H]^+^, [M+Na]^+^, *m/z* 184, N^II*^ | [M+formate]^-^, [M-CH_3_]^-^, phosphocholine |
| **PC** | [M+H]^+^, [M+Na]^+^, *m/z* 184, neutral loss of *sn-*1/*sn-*2 | [M+formate]^-^, [M-CH_3_]^-^, *sn-*1/*sn-*2 RCOO^-^,  loss of *sn-*1/*sn-*2 acyl chain as ketene,  neutral loss of *sn-*1/*sn-*2 RCOOH,  phosphocholine, glycerol-3-phosphate |
| **PC-O** | [M+H]^+^, [M+Na]^+^, *m/z* 184, loss of trimethylamine | [M+formate]^-^, [M-CH_3_]^-^, *sn-*2 RCOO^-^, loss of *sn-*2 acyl chain as ketene, neutral loss of *sn-*2 RCOOH,  glycerol-3-phosphate |
| **PC-P** | [M+H]^+^, [M+Na]^+^, *m/z* 184 | [M+formate]^-^, [M-CH_3_]^-^, *sn-*2 RCOO^-^, loss of *sn-*2 acyl chain as ketene, neutral loss of *sn-*2 RCOOH, phosphocholine |
| **LPC** | [M+H]^+^, [M+Na]^+^, *m/z* 184 | [M+formate]^-^, [M-CH_3_]^-^, *sn-*1 RCOO^-^, loss of *sn-*1 acyl chain as ketene, neutral loss of *sn-*1 RCOOH, phosphocholine, glycerol-3-phosphate |
| **LPC-O** | [M+H]^+^, [M+Na]^+^, *m/z* 184 | [M+formate]^-^, [M-CH_3_]^-^, loss of *sn-*1 acyl chain as ketene, neutral loss of *sn-*1 RCOOH, phosphocholine |
| **LPC-P** | [M+H]^+^, [M+Na]^+^, *m/z* 184 | [M+formate]^-^, [M-CH_3_]^-^, *sn-*1 alkyl, glycerophosphocholine, phosphocholine,  glycerol-3-phosphate |

| **Lipid class** | **Positive-ion ESI-MS** | **Negative-ion ESI-MS** |
| --- | --- | --- |
| **PE** | [M+H]^+^, neutral loss of *m/z* 141 | [M-H]^-^, *sn-*1/*sn-*2 RCOO^-^, loss of *sn-*1/*sn-*2 acyl chain as ketene, neutral loss of *sn-*1/*sn-*2 RCOOH, ethanolamine phosphate, glycerol-3-phosphate |
| **PE-P** | [M+H]^+^, [R_2_COO(C_3_H_5_)OH], [R_1_(CH)_2_HPO_4_(CH_2_)_2_NH_2_],  neutral loss of *m/z* 141 | [M-H]^-^, *sn-*2 RCOO^-^, loss of *sn-*2 acyl chain as ketene, neutral loss of *sn-*2 RCOOH, ethanolamine phosphate |
| **LPE** | [M+H]^+^, neutral loss of *m/z* 141 | [M-H]^-^, *sn-*1 RCOO^-^, loss of *sn-*1 acyl chain as ketene, neutral loss of *sn-*1 RCOOH, ethanolamine phosphate, glycerol-3-phosphate |
| **LPE-P** | [M+H]^+^, neutral loss of *m/z* 141 | [M-H]^-^, neutral loss of plasmenyl group |
| **PI** | n.d. | [M-H]^-^, *sn-*1/*sn-*2 RCOO^-^, loss of *sn-*1/*sn-*2 acyl chain as ketene, neutral loss of *sn-*1/*sn-*2 RCOOH, inositol phosphate - H_2_O, glycerol-3-phosphate |
| **LPI** | n.d. | [M-H]^-^, *sn-*1 RCOO^-^, loss of *sn-*1 acyl chain as ketene, neutral loss of *sn-*1 RCOOH,  inositol phosphate, glycerol-3-phosphate |
| **CE** | [M+NH_4_]^+^, *m/z* 369 | n.d. |
| **Chol** | [M+H-H_2_O]^+^ | n.d. |
| **CAR** | [M+H]^+^, *m/z* 85, *m/z* 144 | n.d. |
| **FA** | n.d. | [M-H]^-^ |
| **MG** | [M+Na]^+^, [M+H-H_2_O]^+^ | n.d. |
| **DG** | [M+Na]^+^, neutral loss of  *sn-*1/*sn-*2, *sn-*1/*sn-*3 fatty acyl | n.d. |
| **TG** | [M+NH_4_]^+^, [M+Na]^+^,  neutral loss of *sn-*1/*sn-*2/*sn-*3 fatty acyl | n.d. |

^*^ The annotation of fragment ions according to Hořejší *et al.* [38] and Merrill *et al.* [39]

Not detected = n.d.
